# Opportunistic binding of EcR to open chromatin drives tissue-specific developmental responses

**DOI:** 10.1101/2022.05.09.491237

**Authors:** Christopher M. Uyehara, Mary Leatham-Jensen, Daniel J. McKay

**Author notes:** Corresponding author: Daniel J. McKay Associate Professor Department of Biology, Department of Genetics, Integrative Program for Biological and Genome Sciences The University of North Carolina at Chapel Hill Chapel Hill, NC 27599, USA.

## Abstract

Steroid hormones perform diverse biological functions in developing and adult animals. However, the mechanistic basis for their tissue specificity remains unclear. In *Drosophila*, the ecdysone steroid hormone is essential for coordinating developmental timing across physically separated tissues. Ecdysone directly impacts genome function through its nuclear receptor, a heterodimer of the EcR and Usp proteins. Ligand binding to EcR triggers a transcriptional cascade, including activation of a set of primary response transcription factors. The hierarchical organization of this pathway has left the direct role of EcR in mediating ecdysone responses obscured. Here, we investigate the role of EcR in controlling tissue-specific ecdysone responses, focusing on two tissues that diverge in their response to rising ecdysone titers: the larval salivary gland, which undergoes programmed destruction, and the wing imaginal disc, which initiates metamorphosis. We find that EcR functions bimodally, with both gene repressive and activating functions, even at the same developmental stage. EcR DNA binding profiles are highly tissue-specific, and transgenic reporter analyses demonstrate that EcR plays a direct role in controlling enhancer activity. Finally, despite a strong correlation between tissue-specific EcR binding and tissue-specific open chromatin, we find that EcR does not control chromatin accessibility at genomic targets. We conclude that EcR contributes extensively to tissue-specific ecdysone responses. However, control over access to its binding sites is subordinated to other transcription factors.

**Significance:** Hormones affect an incredible array of biological processes in both normal development and in disease. In insects, the steroid hormone ecdysone controls processes ranging from neuronal diversification to morphogenesis. Despite its importance, the mechanisms through which ecdysone generates wide-ranging yet tissue-specific responses remain incompletely understood. Like many hormones, ecdysone triggers a cascade of gene expression. At the top of this hierarchy is a nuclear receptor, EcR, which functions both as a hormone receptor and as a transcription factor. However, EcR is not the only transcription factor that functions in the ecdysone cascade; multiple other transcription factors are induced by ecdysone. As a result, the extent to which EcR plays a direct role in regulating tissue-specific responses to ecdysone remains unclear.

## Introduction

Hormones perform an incredible array of biological functions, including regulating physiology (1, 2), reproduction (3, 4), metabolism (5, 6), behavior (6, 7), and development (8, 9). For decades, the insect hormone ecdysone has been studied to delineate the mechanisms by which hormones act on the genome. A key role of ecdysone during development is to serve as a systemic signal to coordinate the timing of events between spatially separated tissues. After release from the prothoracic gland, ecdysone travels through the hemolymph to target tissues where it elicits a response through differential regulation of gene expression. The impact that ecdysone has on target cells is profound. Greater than ten percent of genes respond to ecdysone application in cell culture systems (10). The developmental response to ecdysone is perhaps even more remarkable due to its exquisite specificity, which manifests both spatially and temporally (11). Target tissues can exhibit extremely divergent responses to the same ecdysone pulse. For example, larval structures such as the salivary gland respond to the large pulse of ecdysone that occurs at the end of larval stages by undergoing programmed destruction. By contrast, imaginal tissues such as the wing and leg imaginal discs respond to the late larval ecdysone pulse by initiating the morphogenetic events that create the adult appendages and thorax. Ecdysone also has temporal-specific effects. Pulses of ecdysone occur at stereotypical stages of development, and the same cell lineage responds in distinct ways to different ecdysone pulses. For example, the late larval ecdysone pulse triggers a distinct set of gene expression changes in developing wings relative to those it triggers during the mid-pupal pulse (12). Mechanistically, it is unclear how ecdysone elicits such an extreme breadth of gene expression changes while also maintaining such remarkable spatiotemporal specificity.

In work performed over thirty years ago, ecdysone was found to act on the genome by binding a nuclear receptor heterodimer consisting of EcR and USP (13-15), which bind DNA in a sequence-specific manner through an inverted repeat. Ligand binding to EcR induces structural changes in the receptor that result in differential association with co-activator and co-repressor complexes (16-25). As conceptualized by Ashburner and colleagues through visualization of the puffing program of cultured salivary glands after addition of ecdysone (26), and later brought into molecular detail by multiple labs (27), EcR sits at the top of a transcriptional cascade. Ligand binding to EcR triggers activation of a small set of primary response genes; many of these genes encode transcription factors, which then go on to regulate expression of a larger set of effector genes lower in the cascade. Despite the small number of primary response genes, EcR was recently found to bind extensively across the genome in developing wings, including at many genes with essential roles in wing development (28). Binding of EcR in the wing was also found to be temporally dynamic. These findings extended the canonical Ashburner model by demonstrating that EcR regulates a greater set of target genes beyond those at the top of the ecdysone transcriptional hierarchy. Moreover, they suggested that tissue-specific factors influence EcR binding in the genome, and they raise the important question as to the degree to which EcR autonomously controls binding to its genomic targets.

A major control point in whether transcription factors bind DNA *in vivo* lies at the level of chromatin accessibility (29). Transcription factors and nucleosomes compete for physical access to DNA. Nucleosome-occupancy is often refractory to transcription factor binding, whereas nucleosome-depletion is typically permissive to DNA binding. The importance of chromatin accessibility in controlling access to DNA-encoded information makes identification of the factors that regulate accessibility critical for understanding how cell identities are specified. A link between the ecdysone pathway and regulation of chromatin accessibility was recently uncovered in *Drosophila*. The ecdysone-induced transcription factor E93 was found to specify temporal identity in the wing by controlling accessibility of target enhancers (30). In the absence of E93, late-acting enhancers failed to open and activate, whereas early acting enhancers failed to close and deactivate. Furthermore, precocious expression of E93 in earlier-stage wings resulted in reconfiguration of the larval chromatin accessibility landscape toward pupal wing profiles (31). Thus, the response to ecdysone is mediated at least in part by regulating access to transcriptional enhancers.

In the current study, we investigated the direct role of EcR in controlling tissue-specific ecdysone responses by focusing on two tissues – the larval salivary gland and the wing imaginal disc. We also examined whether EcR, like its primary response gene E93, contributes to tissue-specific ecdysone responses by controlling accessibility of target enhancers. We find that EcR binds extensively across the genome and exhibits a combination of tissue-specific and shared binding profiles. Loss-of-function analysis indicates that EcR is required for the majority of temporal gene expression changes in both tissues. Transgenic reporter analyses demonstrate that binding of EcR is required for target enhancer activity and differential EcR occupancy is associated with tissue-specific enhancer activity. Lastly, although we find that tissue-specific EcR binding is highly correlated with tissue-specific open chromatin, EcR is not required for accessibility of target enhancers. Altogether, our findings support a model in which EcR binds opportunistically to its motif in chromatin made accessible by tissue-specific transcription factors to drive tissue-specific ecdysone responses.

## Results

### Temporal gene expression changes in wings and salivary glands are tissue-specific

In late 3^rd^ instar larvae, rising ecdysone titers initiate morphological and physiological changes that prepare larvae for metamorphosis into adults. Each larval tissue exhibits a distinct transcriptional response during this transition (32). To investigate the role of EcR in mediating tissue-specific gene expression, we focused on two tissues with divergent ecdysone responses: the larval salivary gland and the wing imaginal disc. The larval salivary gland is a secretory organ that has been extensively studied due to the sequence of puffs induced by ecdysone in its polytene chromosomes. Puffs occur at a subset of ecdysone response genes, including many of the early response transcription factors induced by ecdysone, as well as at loci encoding the glue proteins, which are secreted during the larval-to-prepupal transition (33, 34). Shortly after pupariation, the secretory portion of the salivary gland is eliminated via programmed destruction, an event that is dependent on ecdysone signaling (35, 36). By contrast, the wing imaginal disc is not destroyed in response to rising ecdysone titers in third instar larvae. Instead, the disc responds by undergoing a final series of cell divisions, gene patterning events, and cytoskeletal rearrangements to transform from a simple epithelial sheet into a rudimentary progenitor of the adult thorax and wing (37-39) (**Figure 1A**).

**Figure 1.**
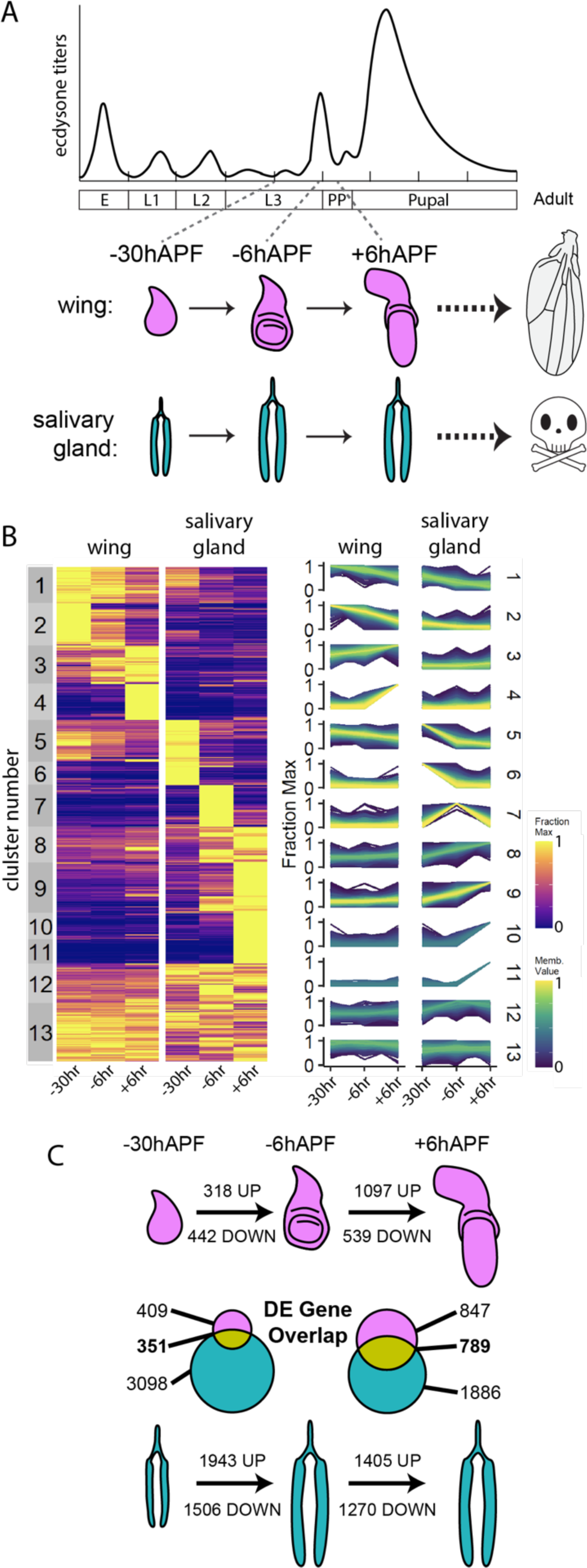
Temporal gene expression profiles in wing and salivary glands are tissue-specific. (A) Schematic of wing and salivary gland development and experimental time points. (B) *Left* Heatmap of RNA-seq values for expressed genes in wild-type wings and salivary glands, following c-means clustering. Values are represented as the fraction of the maximum across all conditions. *Right* Line plots of membership values for genes in each cluster. (C) The number of differentially expressed genes between adjacent timepoints is indicated for each tissue. Venn diagrams depict the overlap of differentially expressed genes between tissues for each time point.

To compare spatiotemporal gene expression changes between the wing and salivary gland, we performed RNA-seq at three time points surrounding the larval-to-prepupal transition: approximately 30-hours prior to the onset of pupariation (–30hAPF), at the wandering stage approximately 6-hours prior to pupariation (–6hAPF), and 6-hours after puparium formation (+6hAPF) (**Figure 1A**). Clustering the RNA-seq data revealed that gene expression profiles are both tissue-specific and temporally dynamic (**Figure 1B**). Although a subset of genes are constitutively expressed in the wing and salivary gland (clusters 12,13), the majority of expressed genes are temporally dynamic in one of the two tissues (clusters 1-11). Observed changes in gene expression are consistent with developmental events occurring at these stages. For example, genes in cluster 2 exhibit high expression levels at –30hAPF that progressively decrease until +6hAPF. This cluster is enriched in GO terms related to mitosis (**Figure S1A, B**), consistent with the rapid proliferation of wing cells during mid-third instar stages that comes to a halt soon after pupariation (40, 41). By contrast, cluster 2 genes are expressed at low levels in salivary glands at all time points, consistent with the lack of a mitotic DNA replication program in larval salivary glands (42). Similarly, genes in clusters 9, 10, and 11, which increase in expression over time in the salivary gland but remain low in wing discs, are enriched in GO terms related to the secretory pathway. This gene expression pattern fits with the production and eventual secretion of glue granules by the salivary gland during pupariation.

To more precisely quantify temporal and tissue-specific gene expression between the wing and salivary gland, we identified differentially expressed genes using DESeq2. Consistent with our clustering results, we found thousands of genes that significantly change in expression across the three time points (**Figure 1C**), with the salivary gland exhibiting a greater number of temporally dynamic genes than the wing. Between –30hAPF and –6hAPF, 760 genes change in the wing and 3,449 genes change in the salivary gland. Between –6hAPF and +6hAPF, 1,636 genes change in the wing, and 2,675 genes change in the salivary gland. A subset of temporally dynamic genes is shared between wings and salivary glands (46%-48% of wing genes, and 10%-29% of salivary gland genes). GO term analysis of these shared genes indicated that metabolism genes are downregulated between –30hAPF and –6hAPF in both wings and salivary glands, whereas genes involved in ecdysone signaling increase in both tissues during this time interval (**Figure S1C**). These gene expression changes are consistent with cessation of feeding behavior triggered by the late-larval ecdysone pulse (43). In sum, gene expression profiles are highly dynamic during the stages leading up to the larval-to-prepupal transition, and the majority of temporally dynamic genes change in a tissue-specific manner.

### Most temporal gene expression changes require ecdysone signaling

Although ecdysone signaling plays an important role in regulating developmental transitions, it is only one of multiple inputs that regulate wing and salivary gland development. Therefore, we next sought to determine the extent to which ecdysone signaling drives developmental progression of wings and salivary glands prior to the larval-to-prepupal transition. During mid-3^rd^ instar, a series of low amplitude ecdysone pulses precede a larger pulse that initiates the onset of pupariation. In the salivary gland, these pulses trigger a puffing cascade, in which a subset of pre-existing intermolt puffs containing the glue genes regress, a small number of puffs are induced at the early response genes, followed several hours later by a larger number of puffs at late-response genes (44). During this same period, the wing undergoes a period of tissue growth and patterning events that spatially define cell identities along the main body axes, including establishing wing versus hinge fates (45) and specifying the position of adult structures such as the wing veins (46). Using a GAL4 driver that is active in both salivary glands and wings, we knocked down EcR via RNAi. Immunofluorescence analysis demonstrated efficient depletion of EcR protein under these experimental conditions (**Figure 2A, B**). At –30hAPF, EcR knockdown has little effect on the morphology of either the wing or salivary gland relative to their wild-type counterparts (**Figure 2A, B**). By contrast, at –6hAPF, *EcR-RNAi* wings are slightly enlarged (28), and *EcR-RNAi* salivary glands are smaller than wild-type glands (**Figure 2B**). In the salivary gland, enlargement of the cytoplasmic lumen is a consequence of the production of glue gene products which are packaged into extracellular vesicles. Glue gene expression is dependent on ecdysone signaling (**Figure 1B**), providing a potential explanation for smaller salivary glands in *EcR-RNAi* animals (47).

**Figure 2.**
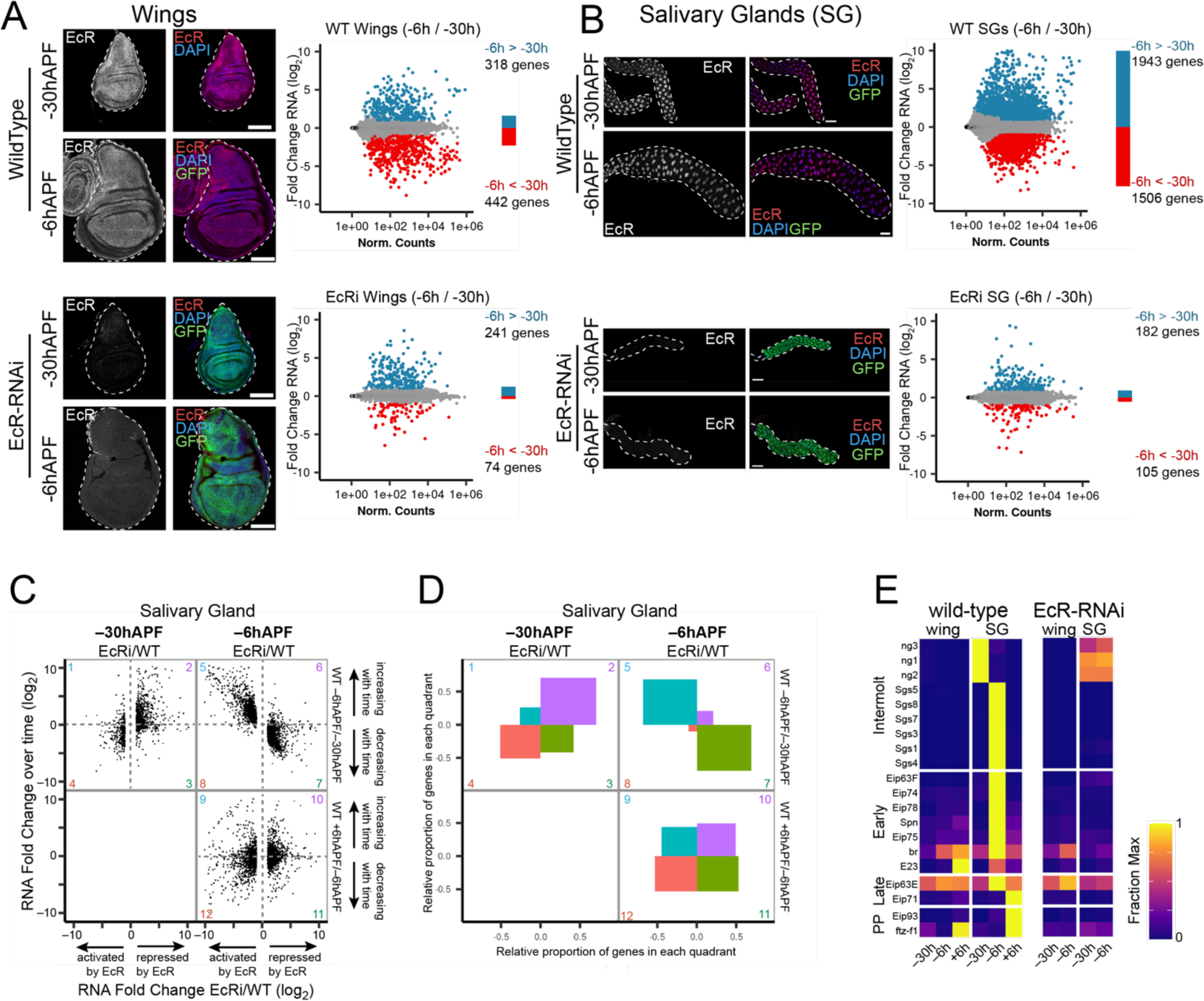
EcR is required to promote genome-wide changes in gene expression over time. (A and B) *Left* Confocal images of wild-type and *EcR-RNAi* wings (A) and salivary glands (B) at –30hAPF and –6hAPF stained with anti-EcR (red) and DAPI (blue). GFP (green) indicates expression of the GAL4 driving RNAi expression. Scale bars are 100μm. *Right* MA plots of RNA-seq data comparing wild-type (*top*) or *EcR-RNAi* (*bottom*) tissues between –30hAPF and –6hAPF. Differentially expressed genes (*P_adj_* < 0.05; absolute log_2_ fold change > 1) are colored red and blue. (C) Scatterplots of RNA-seq values for differentially expressed genes in *EcR-RNAi* salivary glands. The ratio between *EcR-RNAi* and wild-type is shown on the x-axis for –30hAPF and –6hAPF. The ratio between adjacent wild-type stages is shown on the y-axis. (D) Plots indicating the proportion of genes located in each quadrant for the three scatterplots shown in Panel C. (E) Heatmap of RNA-seq values (fraction of max) from wild-type and *EcR-RNAi* tissues for select genes that exhibit ecdysone-dependent puffs in salivary glands. PP indicates prepupal.

To measure the impact of EcR loss of function on gene expression, we performed RNA-seq in *EcR-RNAi* wings and salivary glands at –30hAPF and –6hAPF. We observed a striking failure in temporal gene expression changes in *EcR-RNAi* tissues. In wild-type wing discs, 318 genes increase in expression and 442 genes decrease in expression between –30hAPF and –6hAPF. However, in *EcR-RNAi* wings only 241 genes increase and 74 genes decrease (**Figure 2A**). The gene expression defect was more pronounced in salivary glands. In wild-type glands, 1,943 genes increase in expression and 1,506 gene decrease between –30hAPF and –6hAPF. However, a mere 182 genes increase in expression and 105 genes decrease in *EcR-RNAi* glands (**Figure 2B**). Thus, EcR is required for temporal gene expression changes in late larval tissues. Prior work examining the larval-to-prepupal transition in the wing suggested a bimodal function for EcR, consistent with other ligand-dependent nuclear hormone receptors (48). According to this model, EcR functions primarily as a brake to prevent activation of prepupal genes before the larval-to-pupal transition, whereas following the transition, EcR functions as a trigger to promote expression of prepupal genes (28). To determine if EcR performs a similar role at this earlier developmental stage, we examined the wild-type temporal dynamics of genes differentially expressed in *EcR-RNAi* tissues (**Figure 2C, D**, **Figure S2**). Among the genes that increase between –30hAPF and – 6hAPF in wild-type glands, the great majority require EcR for activation at –6hAPF (**Figure 2C, D** quadrant 5 >> quadrant 6). By contrast, this same set of genes requires EcR for repression at –30hAPF (**Figure 2C, D** quadrant 2 >> quadrant 1). Thus, EcR is required to prevent precocious activation of temporally dynamic genes at –30hAPF, and it is subsequently required for activation of these genes at – 6hAPF, consistent with the brake/trigger mechanism previously proposed in wings. However, a brake/trigger model only partially explains EcR’s role in temporal gene regulation. Among EcR target genes that decrease between –30hAPF and –6hAPF in wild-type glands, most require EcR for repression at –6hAPF (**Figure 2** **D, E** quadrant 7 >> quadrant 8). By contrast, this same set of genes requires EcR for activation at –30hAPF (**Figure 2D, E** quadrant 4 > quadrant 3). Thus, EcR also switches from an activator to a repressor between –30hAPF and –6hAPF. This finding challenges a simple model of EcR switching from a repressor to an activator as ecdysone titers rise in late larvae. A similar signature is also apparent in the wing, where EcR target genes that increase between –30hAPF and –6hAPF switch from requiring EcR for repression at –30hAPF to requiring EcR for activation at –6hAPF (**Figure S2**). Likewise, EcR target genes that decrease between –30hAPF and –6hAPF in wild-type wings switch from requiring EcR for activation at –30hAPF to requiring EcR for repression at –6hAPF. Interestingly, this signature role of EcR as a switch continues in the wing during the larval-to-prepupal transition (**Figure S2** quadrants 9-12), but the relationship breaks down in salivary glands during the larval-to-prepupal transition (**Figure 2D, E** quadrants 9-12), suggesting tissue-specific roles of EcR in mediating responses to the third larval ecdysone pulses.

### Expression of ecdysone-induced transcription factors is tissue-specific

Examination of puffing profiles in the salivary gland has uncovered a transcriptional cascade that unfolds in response to rising ecdysone titers during third instar stages. Included in this cascade is a set of transcription factors that mediate ecdysone-induced gene expression responses in the salivary gland and a wide range of other tissues. Given the differential impact of EcR loss of function on temporal gene expression profiles in wings and salivary glands, we investigated whether these ecdysone-induced transcription factors also diverge in expression between wings and salivary glands. We compiled a list of ecdysone-induced transcription factors plus an additional set of genes that exhibit clearly defined puffing profiles in salivary glands (27, 49–56). This list was divided into four categories based on the timing of their puffing profiles: 1) intermolt puffs, containing the glue genes, which regress with rising ecdysone titers; 2) early puffs, which are rapidly induced with rising ecdysone titers; 3) late puffs, which are induced several hours after early puffs; and 4) prepupal puffs, which are induced after the onset of pupariation. Our RNA-seq data mirror the temporal progression of the puffing cascade in wild-type salivary glands (**Figure 2E**), with genes residing in intermolt puffs expressed earliest, followed by early, late, and prepupal genes. In the wing, the early and late-response transcription factors *broad* (*br*) and *ftz-f1*exhibit similar temporal profiles relative to the salivary gland. However, other ecdysone-induced transcription factors such as *E74*, *E75*, and *E78* are not induced to high levels during this time interval in the wing. Notably, these transcription factors are highly induced in the wing later during pupal stages (**Figure S3**) (40). Thus, the same set of ecdysone-induced transcription factors are employed in both wings and salivary glands; however, the timing of their maximal deployment varies considerably between tissues. Consistent with the role of these transcription factors as primary response ecdysone target genes, each of them failed to activate in EcR-RNAi salivary glands (**Figure 2E**). Surprisingly, the only ecdysone-induced transcription factor expressed in wild-type wings in late larvae, *br*, was still activated despite EcR loss of function. *Br* was modestly activated in EcR-RNAi salivary glands, albeit at lower levels than in wild-type glands, and in both tissues, *br* expression levels increased over time despite EcR loss of function. Thus, other temporal inputs besides ecdysone signaling likely regulate *br* expression in both salivary glands and wings.

### EcR exhibits a complex mixture of tissue-specific and shared DNA binding profiles

As described above, transcriptional responses to ecdysone are hierarchically regulated; EcR/Usp induces expression of primary response transcription factors such as *br* and *E93*, which contribute to regulation of downstream effector genes. This hierarchical regulatory structure has left the direct role of EcR in mediating ecdysone responses obscured. To examine the potential function of EcR in directing tissue-specific responses to ecdysone, we performed CUT&RUN on –6hAPF wing imaginal discs and salivary glands. We found that EcR exhibits both tissue-specific and shared DNA binding profiles (**Figure 3**). We used DESeq2 to define three categories of EcR binding: wing-enriched, salivary gland-enriched, and shared (no significant difference between tissues). We observed a mixture of wing-enriched binding, salivary gland-enriched binding, and shared binding, even at the level of individual target genes (**Figure 3A, B**). Notably, tissue-specific EcR binding is associated with tissue-specific target gene expression (**Figure 3C**). Genes that have wing-enriched EcR binding sites exhibit higher expression in wing imaginal discs relative to salivary glands. Likewise, genes that have salivary gland-enriched binding sites are more highly expressed in salivary glands relative to wings (**Figure 3C**). Moreover, EcR binding is associated with differential target gene expression upon EcR loss of function (**Figure 3D**). Genes bound by EcR in the wing imaginal discs tend to be upregulated in *EcR-RNAi* wings, particularly at – 30hAPF. Similarly, genes bound by EcR in the salivary gland tend to be upregulated in *EcR-RNAi* glands at –30hAPF. Notably, this relationship is inverted at –6hAPF; genes bound by EcR in the salivary gland tend to be downregulated upon EcR loss of function at –6hAPF. These findings are consistent with EcR functioning as both a repressor and an activator, and they suggest that at least some of the observed changes in gene expression in *EcR-RNAi* tissues are due to direct binding by EcR to target genes.

**Figure 3.**
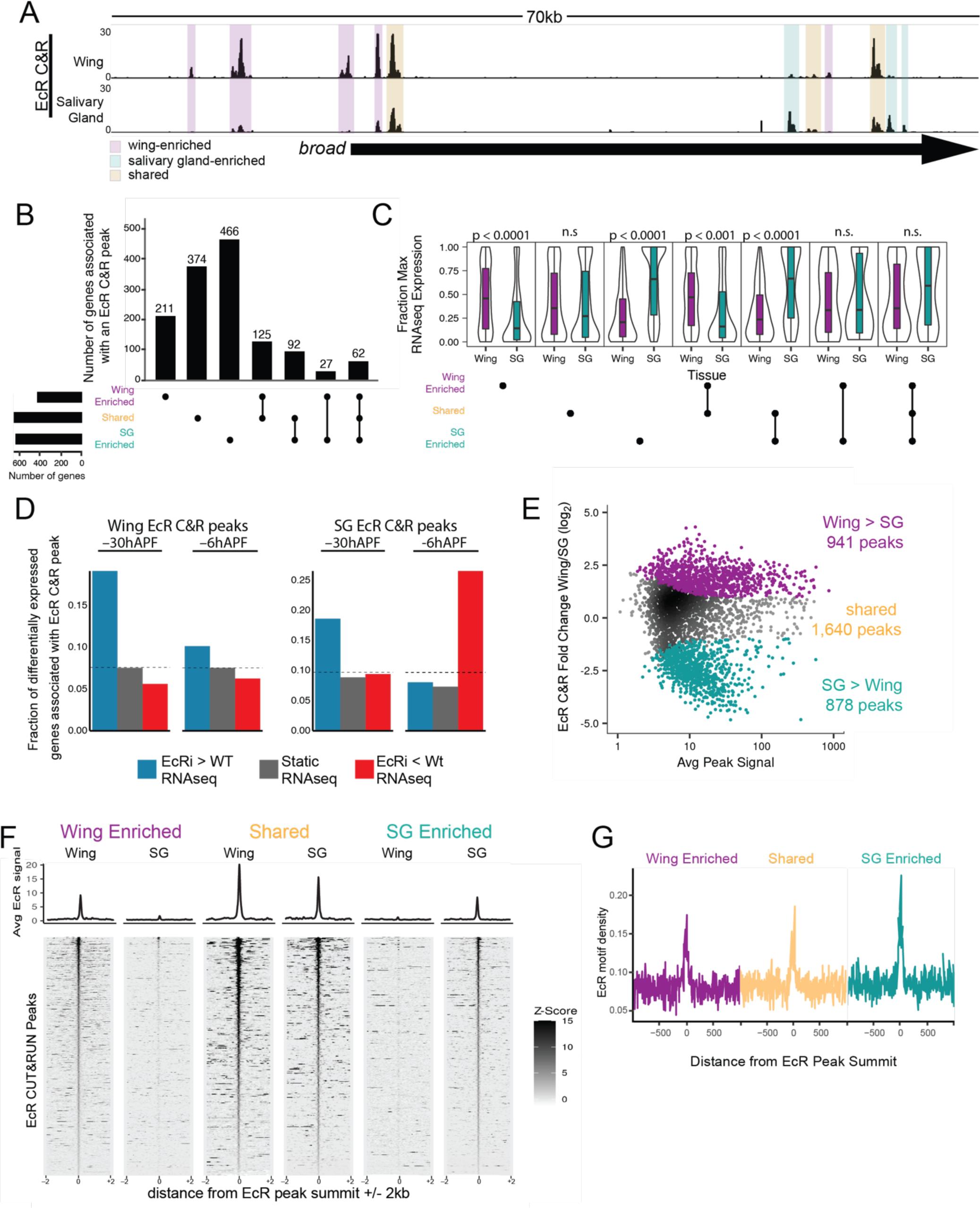
EcR binding is tissue-specific. (A) Browser shot of EcR CUT&RUN data from –6hAPF wings and salivary glands, with examples of wing-enriched, salivary gland enriched, and shared peaks highlighted by colored boxes. (B) Upset plot of the number of genes associated with at least one EcR CUT&RUN peak in each tissue, split by EcR peak category. Each EcR-bound gene is only represented once. (C) Violin plots of RNA-seq values (fraction of max) of genes bound by EcR in at least one tissue. P values represent comparisons between wing and salivary gland for each upset category (Wilcoxon rank sum test with Bonferroni correction). (D) Bar plot of the fraction of differentially expressed EcR-bound genes in *EcR-RNAi* wings and salivary glands. The dotted lines indicate the fraction of all expressed genes in each EcR peak category. (E) MA plot depicting –6hAPF EcR wing and salivary CUT&RUN values in a union set of EcR CUT&RUN peaks. Differential peaks are indicated with colored dots. (F) Heatmaps and average signal plots of EcR CUT&RUN signal for wing-enriched, shared, and salivary gland-enriched peaks centered on peak summits. For shared peaks, the wing summit was used. (G) Line plots (10bp bins) of EcR motif density +/-1kb from EcR peak summits for the three EcR binding categories.

At the genome-wide level, approximately two-thirds of each tissue’s EcR binding sites are shared between wings and salivary glands, and one-third is specific to each tissue (941 wing-enriched EcR peaks, 878 salivary gland-enriched EcR peaks, 1,640 shared EcR peaks) (**Figure 3E, F**). Relative to the whole genome, EcR binding is enriched in promoters and introns (**Figure S4A**). EcR binding sites also tend to be clustered throughout the genome, with approximately half of EcR peaks found within 5kb of another peak (**Figure S4B, C**). Primary response ecdysone target genes are particularly enriched EcR binding peaks (**Figure S5**). To determine whether differences in DNA sequence composition contributes to differential EcR binding between wings and salivary glands, we examined EcR motif content for the three categories of EcR peaks. We observed no difference in the quality or density of EcR DNA binding motifs in wing-enriched, salivary gland-enriched, or shared EcR peaks (**Figure 3G, S4E, F**), indicating that EcR motifs do not drive global differences in EcR DNA binding profiles between wings and salivary glands.

### Tissue-specific binding is associated with tissue-specific open chromatin

The mixture of tissue-specific and shared EcR binding sites at target genes indicates that the ecdysone response is supported by a complex *cis*-regulatory architecture. To gain insight into the function of genomic regions bound by EcR, we performed FAIRE-seq on wings and salivary glands at –6hAPF (**Figure 4A**). FAIRE enriches for regions of DNA that are locally depleted of nucleosomes, thus it can be used as a proxy for identifying *cis*-regulatory elements genome wide (57). We observed tissue-specific open chromatin profiles in wings and salivary glands, with many differentially accessible sites found in each tissue, as well as a set of FAIRE peaks that are shared between the two tissues. In total, we identified 3,697 peaks that are more accessible in wings, 4,406 peaks that are more accessible in salivary glands, and 1,856 peaks that are not differentially accessible (i.e. “shared”) (**Figure 4B**). These differences in FAIRE signal contrast with more similar chromatin accessibility profiles observed between wing, leg, and haltere imaginal discs at the same developmental stage (58). Notably, differential accessibility between wings and salivary glands is associated with differential gene expression, with wing-enriched FAIRE peaks associated with higher gene expression in wings, and salivary gland-enriched FAIRE peaks associated with higher gene expression in salivary glands (**Figure 4C**). To examine the relationship between chromatin accessibility and EcR binding, we calculated the difference in FAIRE-seq signal between wings and salivary glands at EcR CUT&RUN peaks. We found that wing-enriched EcR peaks are more accessible in the wing than in the salivary gland, while salivary gland-enriched peaks are more accessible in the salivary gland (**Figure 4D**). Thus, tissue-specific EcR binding is associated with tissue-specific chromatin accessibility.

**Figure 4.**
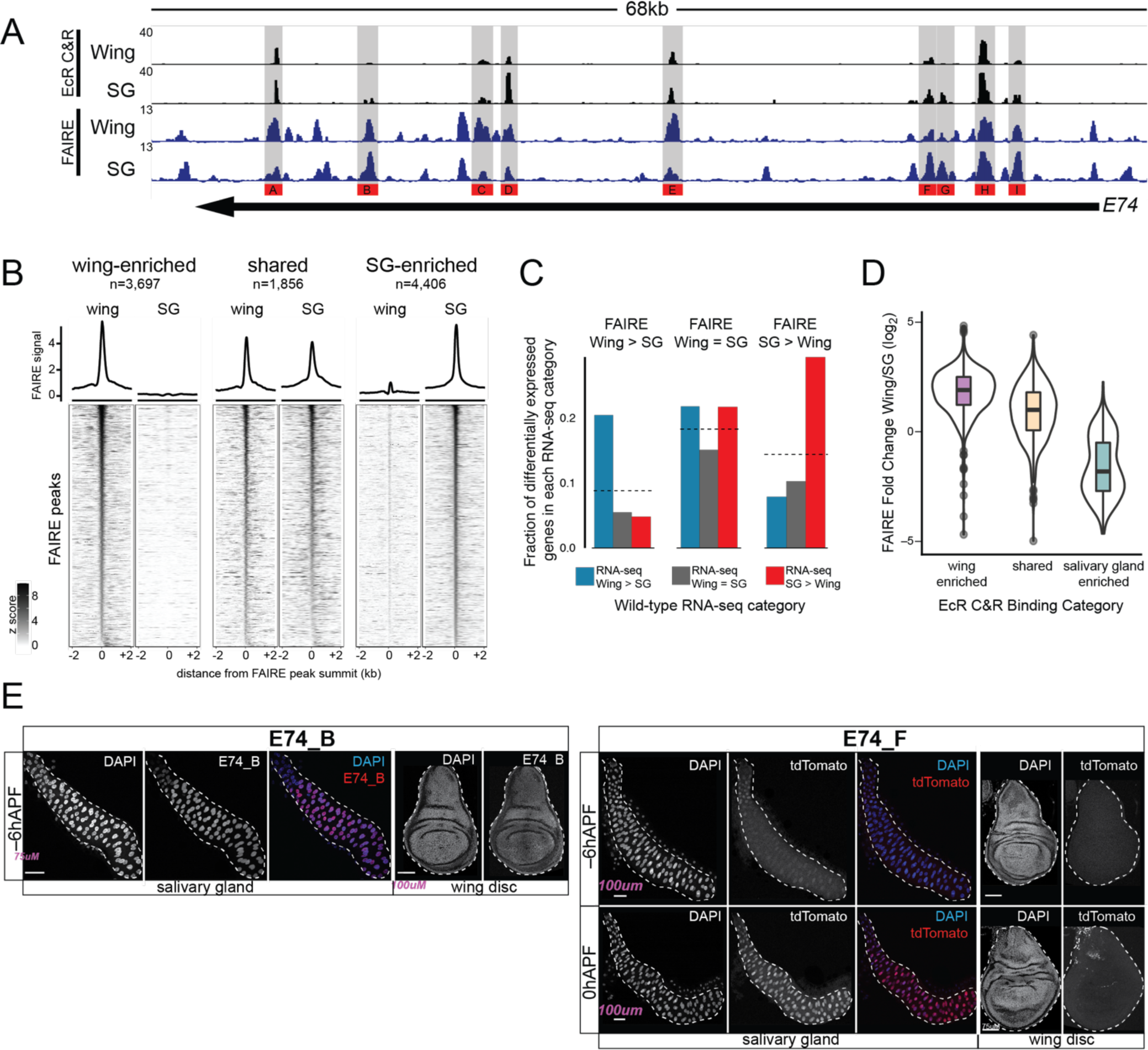
Tissue-specific EcR binding is associated with tissue-specific open chromatin and enhancer activity. (A) Brower shot of EcR CUT&RUN and FAIRE data from –6hAPF wings and salivary glands. Regions cloned for transgenic reporter analysis are indicated. (B) Heatmaps and average signal plots of FAIRE signal surrounding wing-enriched, shared, and salivary gland-enriched FAIRE peaks centered on the peak summit. For shared peaks, the wing summit was used. (C) Bar plots indicating the fraction of differentially expressed genes between wild-type wings and salivary glands that overlap a FAIRE peak in each of the three indicated FAIRE categories. Dotted lines indicate the fraction of all expressed genes that overlap a FAIRE peak in each category. (D) Violin plots of ratio of FAIRE signal between wings and salivary glands for the three indicated EcR CUT&RUN peak categories. (E) Confocal images of *E74_F* (*top*) and *E74_B* (*bottom*) enhancer activity in salivary glands (*left*) and wings (*right*). –6hAPF and 0hAPF are shown for *E74_F*. –6hAPF is shown for *E74_B*. tdTomato signal for *E74_F* at 0hAPF is derived from cells associated with the trachea; the enhancer is inactive in the main epithelium.

To examine the *cis*-regulatory potential of EcR-bound regions, we turned to transgenic reporter assays. We cloned nine EcR-bound regions from the primary-response gene, E74. We also evaluated an EcR bound region that had been previously characterized in the wing imaginal disc (*br^disc^*), as well as an EcR bound region that overlaps a transgenic reporter generated by the Janelia FlyLight collective (*GMR79E07*). These cloned regions exhibit a range of relative and absolute values for EcR binding amplitude and degree of chromatin accessibility between –6hAPF wings and salivary glands. In total, ten out of eleven tested regions act as transcriptional enhancers. The *br^disc^*enhancer, which is strongly enriched for accessibility and EcR binding in wings relative to glands, is active throughout the wing but only in a small number of cells in the salivary gland, corresponding to the duct and imaginal ring cells (**Figure S6A**). Conversely, the *GMR79E07* region, which is accessible and EcR bound only in the salivary gland, is active only in glands and not in the wing (**Figure S6A**). Interestingly, the *E74_F* enhancer, which is bound by EcR in both the salivary gland and wing and is enriched in accessibility in the salivary gland, exhibits salivary gland-specific enhancer activity that increases dramatically from – 6hAPF to +6hAPF (**Figure 4E**). The *E74_D* and *E74_G* enhancers are also temporally dynamic, increasing in activity in late 3^rd^ instar larvae and prepupae (**Figure S6**). The *E74_A*, *E74_B*, and *E74_E* enhancers, which exhibit similar relative levels of EcR binding between wings and glands, but which differ in the absolute levels of EcR binding between the enhancers, are transcriptionally active in both tissues at –6hAPF (**Figure 4E, S6**). The *E74_H* enhancer is active at low levels only in the transitional cells of the salivary gland but is inactive in wings. The *E74_I* region did not exhibit enhancer activity in either tissue despite its accessibility and EcR occupancy in both tissues. We conclude that many EcR binding sites correspond to functional enhancers; further, relative differences in accessibility and amplitude of EcR CUT&RUN signal are associated with differential enhancer activity. However, absolute levels of accessibility or EcR CUT&RUN signal are poor predictors of the strength or pattern of enhancer activity.

### EcR knockdown does not cause global changes in chromatin accessibility

The experiments described above indicate that EcR can bind enhancers that exhibit tissue-specific activity, suggesting that EcR plays a direct role in controlling tissue-specific responses to ecdysone. Moreover, at the genome-wide level, tissue-specific EcR binding is closely associated with tissue-specific open chromatin, raising the possibility that EcR directs each tissue’s response to ecdysone by controlling accessibility of tissue-specific target enhancers. To test this hypothesis, we performed FAIRE-seq in *EcR-RNAi* wings at –6hAPF. If EcR controls chromatin accessibility, then EcR-bound sites are predicted to decrease in accessibility in *EcR-RNAi* wings. Alternatively, if EcR is not required to open chromatin, then accessibility of EcR-bound sites is predicted to remain unaffected. Comparison of wild-type and *EcR-RNAi* wing FAIRE-seq profiles revealed few differences in open chromatin (**Figure 5A, B**). Examination of FAIRE-seq signal specifically at EcR CUT&RUN peaks likewise revealed no global changes in chromatin accessibility in *EcR-RNAi* wings (**Figure 5C**). In total, we observed only 132 FAIRE peaks that are more accessible in EcR-RNAi relative to wild-type and only 150 peaks that are less accessible in EcR-RNAi wings relative to wild-type. To put these numbers into perspective, wings and salivary glands collectively exhibit 1,706 tissue-specific EcR peaks and 4,556 tissue-specific FAIRE peaks. To investigate if EcR knockdown causes more subtle effects on chromatin accessibility, we examined FAIRE-seq signal at FAIRE peaks separated into two categories depending on whether they overlap an EcR CUT&RUN peak. We observed a slight decrease in accessibility in *EcR-RNAi* wings for EcR-bound FAIRE peaks relative to unbound FAIRE peaks (**Figure 5D**). Collectively, these data are consistent with a model in which EcR does not function as a pioneer factor to control chromatin accessibility at target enhancers. Binding of EcR may subtly contribute to the amplitude of accessibility at its targets; however, it does not function as a binary switch to open target site chromatin.

**Figure 5.**
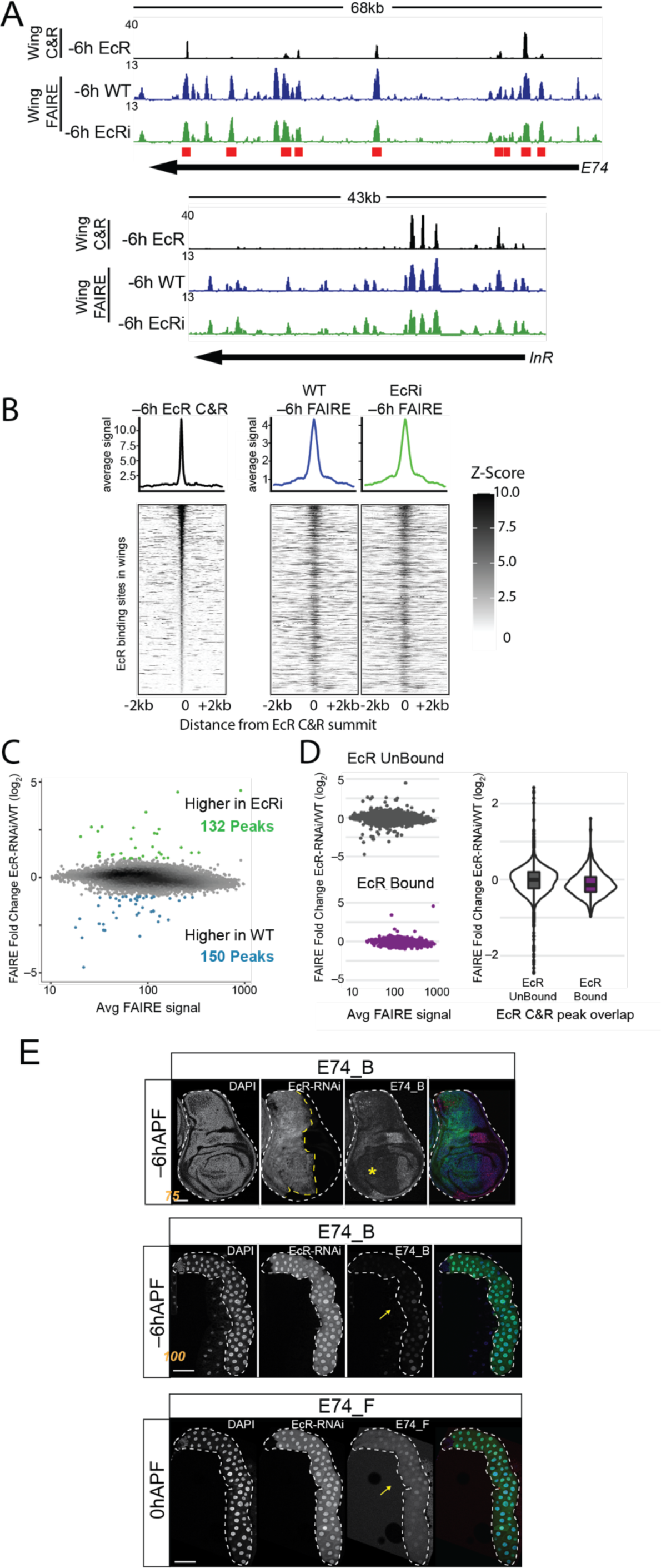
EcR loss of function has minimal impact on chromatin accessibility. (A) Browser shots of EcR CUT&RUN signal from –6hAPF wild-type wings, and FAIRE signal from –6hAPF wild-type and *EcR-RNAi* wings. Red boxes indicate the location of cloned regions from the *E74* locus. (B) Heat maps and average signal plots of FAIRE signal in –6hAPF EcR CUT&RUN peaks from –6hAPF wild-type and *EcR-RNAi* wings. EcR CUT&RUN signal is shown on the left for reference. (C) MA plot of FAIRE signal in the union set of FAIRE peaks from –6hAPF wild-type and *EcR-RNAi* wings. (D) MA plots (*left*) and violin plot (*right*) of the FAIRE data shown in panel C separated according to their overlap with an EcR CUT&RUN peak. (E) Confocal images of *E74_B* activity (red) in –6hAPF wings (*top*) and salivary glands (*middle*), and *E74_F* activity in 0hAPF salivary glands (*bottom*). Cells expressing EcR-RNAi are marked with GFP (green). The yellow asterisk and arrow indicate loss of enhancer activity in wings and glands, respectively.

To evaluate EcR’s role in regulating individual target enhancers, we examined the activity of E74 transgenic reporters upon EcR loss of function. In all cases tested, we found that EcR is required for target enhancer activation in both wings and salivary glands without affecting enhancer accessibility (**Figure 5**, **Figure S6**). Using *Ci-GAL4* to drive *EcR-RNAi* in the anterior compartment of the wing imaginal disc, we observed loss of *E74_B* activity (**Figure 5E**, **top**). Likewise, EcR knockdown in the salivary gland resulted in loss of *E74_B* activity (**Figure 5E, middle**). We obtained similar results with other *E74* enhancers active in both wings and salivary (**Figure S6**). Notably, despite marked differences in the amplitude of EcR CUT&RUN signal at target enhancers such as *E74_E* and *E74_B* enhancer, both enhancers fully depend on EcR for activation, indicating that even low-amplitude binding can be biologically relevant. Temporally dynamic enhancers such as *E74_D*, *E74_G*, and *E74_F* also depend on EcR for activation during the larval-to-prepupal transition (**Figure 5E, S6**). We also tested whether binding of EcR is required for repression of enhancers such as *E74_D*, which is bound by EcR in the wing, where it is inactive. However, knockdown of EcR did not affect *E74_D* activity in –6hAPF wings (**Figure S6**), indicating that EcR can bind targets without having an obvious role in transcriptional regulation. As noted above, knockdown of EcR throughout the wing imaginal disc did not affect chromatin accessibility at any of these target enhancers (**Figure 5A**). Thus, despite being required for enhancer activation, EcR does not control accessibility of its genomic targets.

### EcR binding profiles are shared between wing and leg imaginal discs

Thus far, our findings support a model in which different developmental responses to ecdysone involve direct binding of EcR to both shared and tissue-specific enhancers. However, despite a strong association between EcR binding and tissue-specific open chromatin, EcR is not required for chromatin accessibility at its targets. We reasoned that this pattern of EcR binding could result from differential recruitment of EcR to target enhancers by tissue-specific transcription factors. To further investigate the forces influencing EcR binding site selectivity, we turned to the leg imaginal disc. The distinct identities of wing and leg imaginal discs are driven by differential expression of master regulator transcription factors. For example, *vestigial*, *apterous*, and the *Iroquois complex* transcription factors specify wing identity, whereas *Sp1*, *buttonhead*, and *Distalless* specify leg identity. Despite differential expression of master regulator transcription factors, and differences in their DNA binding profiles between imaginal discs, wing and leg imaginal discs have exceptionally similar open chromatin profiles (58, 59). If the master regulators of tissue identity recruit EcR to target enhancers, then it is predicted that EcR binding profiles would differ between wing and leg imaginal discs.

Consistent with prior findings, the open chromatin profiles of –6hAPF wing and leg imaginal discs are highly similar. By contrast, the number of differential FAIRE peaks between salivary glands and wings or legs is an order of magnitude greater (**Figure 6A, B**, **4B**, **Figure S7**). Comparison of EcR CUT&RUN data from wings and legs revealed strikingly similar DNA binding profiles: only 36 EcR peaks were identified as differentially bound between wing and leg imaginal discs, as compared to 2,432 EcR peaks identified as shared between the two tissues (**Figure 6C**). Accordingly, EcR binding profiles in wing and leg are far more similar to each other than either is to the salivary gland (**Figure 6B**). The small number of differentially bound EcR peaks between wing and leg imaginal discs correspond to the relatively small number of differentially accessible FAIRE peaks between the two tissues (**Figure S7**). The similarity between EcR binding profiles in the wing and leg argues against a primary role of tissue-specific transcription factors guiding EcR to target enhancers. Instead, the high degree of overlapping peaks between wings and legs supports a model in which EcR opportunistically binds its motif in chromatin already made accessible by other transcription factors (**Figure 6D**).

**Figure 6.**
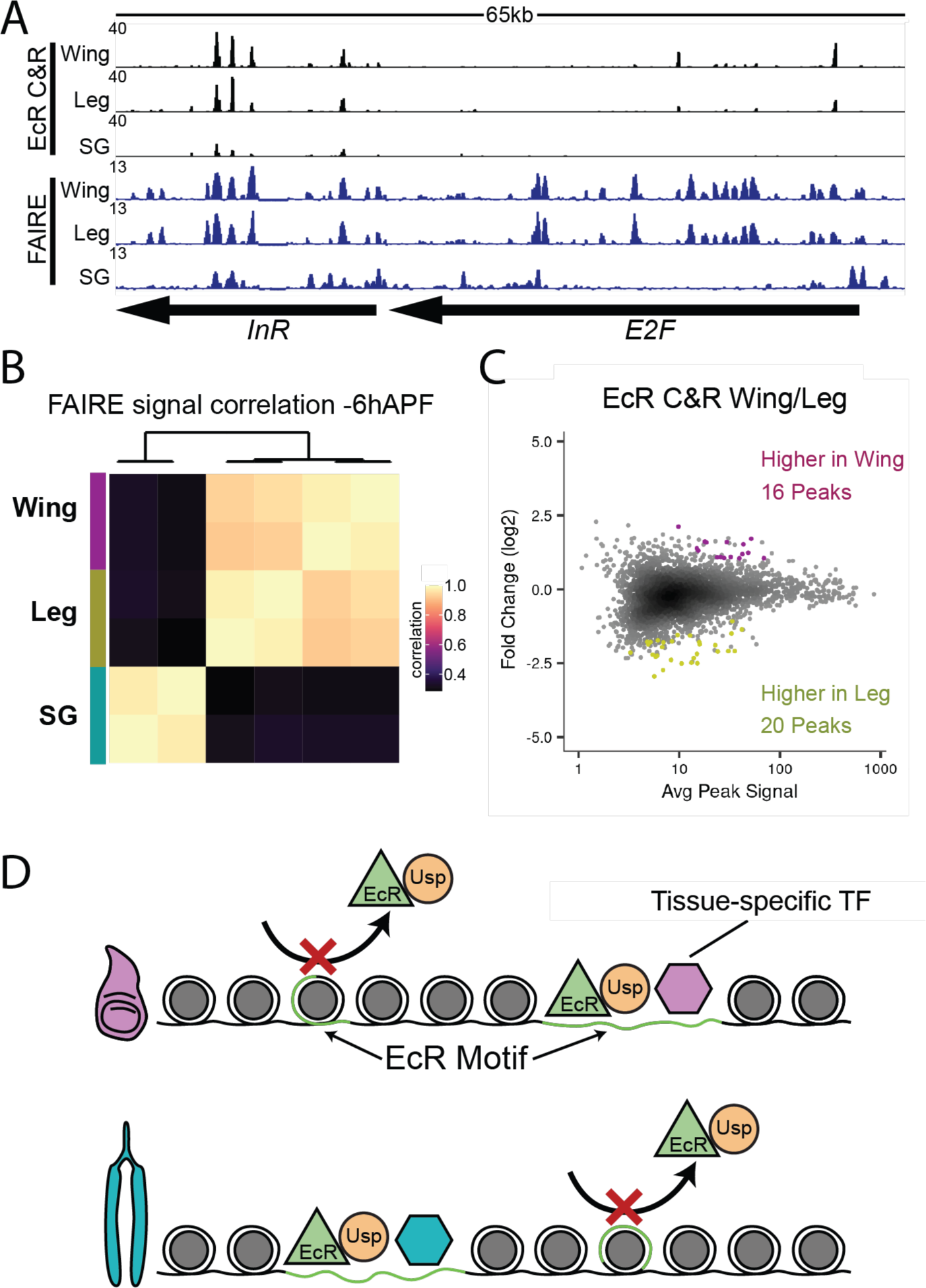
EcR binding profiles in legs and wings are more similar than those in salivary glands. (A) Browser shot of EcR CUT&RUN and FAIRE signal from –6hAPF wild-type wings, legs, and salivary glands. (B) Heatmap of the Pearson correlation coefficient for EcR CUT&RUN signal between tissues. (C) MA plot of EcR CUT&RUN signal between –6hAPF wild-type wings and legs. Differential peaks are indicated by colored dots. (D) Model depicting the dependence of EcR (green triangle) on tissue-specific transcription factors (hexagons) for DNA binding.

## Discussion

The response to ecdysone is both wide-ranging and transcriptionally diverse; tissues throughout the animal respond to the same pulse of ecdysone in markedly different ways. How this specificity is achieved remains incompletely understood. In this work, we find that tissue-specific ecdysone responses are due at least in part to differential binding of EcR to tissue-specific enhancers. Although EcR is required for activation of target enhancers, it does not control their accessibility. Thus, despite a direct and widespread role in controlling tissue-specific responses to ecdysone, EcR remains subordinate to chromatin accessibility programs determined by other transcription factors.

### Tissue- and temporal-specific gene expression profiles in the wing and salivary gland

The last thirty-six hours of larval development marks a period of incredible change in *Drosophila* development; both larval and imaginal tissues must rapidly and coordinately complete developmental programs required for the impending transformation of the body plan during pupal stages. Salivary glands begin producing and storing glue gene products, prior to secreting them during pupariation. Soon thereafter, the larval portion of the gland undergoes programmed destruction (60-63). By contrast, third-instar wing imaginal discs continue extensive tissue patterning and growth, and then begin the process of metamorphosis into an adult appendage (64, 65). Consistent with these developmental changes, our RNA-seq time course identified a substantial number of differentially expressed genes. Although a subset of temporal gene expression changes are shared between wings and salivary glands, most changes are tissue specific. Unexpectedly, we identified many more differentially expressed genes in the salivary gland than in the wing. One possible explanation for this skew is differences in cell-type heterogeneity between the two tissues. Salivary glands are composed primarily of three cell types: secretory cells, duct cells, and imaginal ring cells (66). Due to the shared function of any one of these cell types, it is likely that their transcriptional profiles are highly similar, making RNA-seq performed on whole glands particularly sensitive to gene expression changes. By contrast, the wing imaginal disc exhibits more cell-type heterogeneity (67), meaning that a smaller fraction of cells across the tissue share transcriptional profiles, thus decreasing the sensitivity of RNA-seq to detect gene expression changes from whole wing imaginal discs. As a result, we are likely underestimating the number of temporal gene expression changes on a cell-by-cell basis in wings relative to salivary glands. Despite different sensitivities in quantifying transcriptional profiles, the developmental impact of EcR loss of function is more pronounced in the salivary gland relative to the wing. Morphology of *EcR-RNAi* glands is more disrupted than wings at –6hAPF, and a greater number of genes are deregulated in *EcR-RNAi* glands relative to *EcR-RNAi* wings (**Figure 2A, B**). These findings suggest that salivary glands depend on ecdysone signaling more heavily than wings. This interpretation is consistent with the inability of the EcR ligand binding domain to activate transcription in late third instar imaginal discs despite robust activity in larval tissues such as the salivary gland (68). It is unclear why larval and imaginal tissues would exhibit differential responsiveness to the late-larval ecdysone pulse. One possibility is that salivary gland and wing imaginal disc cells differ in their ability to transport ecdysone across the cell membrane. A membrane transporter was recently identified that is required for cellular import of ecdysone, challenging the prevailing model that ecdysone enters cells through simple diffusion (69, 70). However, the gene encoding this transporter, *Oatp74D*, is expressed at high levels in both wings and salivary glands (**Table S1**), suggesting that ecdysone import should be similar in both tissues. It is also possible that salivary glands and wings differ in their ability to convert ecdysone to its physiologically active form, 20-hydroxyecdysone. Consistent with this hypothesis, the gene encoding the enzyme responsible for this conversion, *shade*, is expressed more highly in salivary glands than in wings (**Table S1**). However, *shade* has been shown to function in a tissue non-autonomous manner (71), making it unclear whether differential *shade* expression is sufficient to explain differences in ecdysone responsiveness between wings and salivary glands. Lastly, it is possible that EcR gene regulatory complexes differ between the wing and salivary gland. EcR interacts with both transcriptional co-repressors (72-74) and co-activators (75-81). The composition or relative abundance of these complexes may differ between wings and glands, resulting in distinct transcriptional outputs despite similar ecdysteroid levels.

Regardless of any differences in the impact of ecdysone signaling on salivary glands and wings, EcR is clearly required for the majority of gene expression changes between –30hAPF and –6hAPF in both tissues, indicating that the ecdysone pathway is the primary temporal gene expression driver for both tissues at this time. Rising ecdysone titers at the end of larval development are likely responsible for triggering EcR-dependent gene expression changes. Like other nuclear receptors, EcR is thought to function as a hormone-dependent switch (48). Along these lines, prior work from our lab provided support for a brake/trigger model of EcR action (28). We found that EcR is required to prevent precocious activation of prepupal genes in the larval wing (the brake), and it is subsequently required for activation of these genes in prepupal wings (the trigger). This is consistent with a model in which EcR switches from a repressor to an activator with increasing hormone titers. Here, we observed similar effects in salivary glands and wings; genes that increase with developmental time become precociously active and subsequently fail to reach wild-type levels in the absence of EcR (**Figure 2C, D**). However, the additional RNA-seq data from –30hAPF reveal that this model is incomplete. Loss of EcR function not only results in precocious activation of genes that increase with time, but it also results in precocious repression of genes that decrease with time, suggesting that EcR switches from an activator to a repressor over the same time interval that it switches from a repressor to an activator. While many of these defects in gene expression are likely to be secondary consequences due to the requirement of EcR in activating primary response transcription factors (**Figure 2E**), we also provide evidence for a direct role of EcR in target gene regulation. Most *E74* enhancers we evaluated depend on EcR for activation, whereas the *br^disc^* enhancer is repressed by EcR (28). Moreover, at the genome-wide level, EcR binding sites are enriched in genes that are repressed by EcR in both wings and salivary glands (**Figure 3D**). These findings argue against simple models in which rising hormone titers trigger a switch in EcR’s interacting partners from co-repressors to co-activators. Instead, they suggest that hormone binding can also induce EcR to associate with co-repressor complexes. Indeed, the co-repressor Mi2 has been shown to interact with EcR in an ecdysone-dependent manner (72, 82). The data provided here suggest that hormone-induced repression is pervasive.

### A complex *cis*-regulatory architecture supports tissue-specific ecdysone responses

The possibility that EcR binding might vary across tissues was first suggested by observations that the puffing cascade differs between the larval fat body and salivary gland (56). Since then, multiple cis-regulatory elements with functional EcR binding sites have been identified. Several of these enhancers control tissue-specific gene expression (83-85). Our EcR DNA binding profiles in wings and salivary glands both validate and extend these findings. We find that EcR binds extensively across the genome. In fact, most EcR binding sites reside at genes that are not conventionally defined as primary response genes, indicating that EcR directly regulates genes located in both the upper and lower tiers of the ecdysone-response hierarchy. Moreover, one-third of EcR binding sites are specific to either wings or salivary glands, suggesting that a significant portion of tissue-specific responses to ecdysone are driven through direct EcR binding. Surprisingly, tissue-specific EcR binding occurs not only at terminal effector genes that mediate tissue-specific responses (such as those involved in glue secretion in the salivary gland), but also at many primary response genes. A notable example of this complex *cis*-regulatory architecture is found at the *broad* locus, a well-characterized primary response gene. Even though *broad* plays a central role in mediating the response to ecdysone in many tissues, our analyses reveal tissue-specific differences in its regulation. RNA-seq profiles indicate that the timing of *broad* expression varies between the wing and salivary gland. Peak *broad* expression is observed at –6hAPF in the salivary gland, whereas in the wing, *broad* peaks at +6hAPF. Moreover, ecdysone-dependent regulation of *broad* differs between the two tissues. *Broad* expression is strongly reduced upon EcR loss of function in the salivary gland, but its activation occurs nearly normally in *EcR-RNAi* wings. We find that differential *broad* regulation stems from differential binding of EcR to enhancers with tissue-specific activity (**Figure 3, 4**).

A second primary response gene, *E74*, also exhibits markedly distinct expression profiles between wings and salivary glands driven by differential EcR binding. Like *broad*, *E74* is expressed in both wings and salivary glands at all three time points measured. *E74* expression increases ∼17-fold in the salivary gland to a peak at –6hAPF, before decreasing ∼60-fold to a nadir at +6hAPF (**Figure S2, Table S1**). In the wing, *E74* expression increases only ∼1.4-fold from –30hAPF to –6hAPF, before decreasing back to its prior levels at +6hAPF. EcR loss of function reveals that increasing *E74* expression at –6hAPF is dependent on ecdysone in both wings and salivary glands. However, a more extensive RNA-seq time course indicates that the true peak of *E74* expression in the wing does not come until +44hAPF (**Figure S2**). Transgenic reporter analyses reveal that tissue-specific differences in *E74* expression derive from tissue-specific differences in enhancer activity; *E74_D*, *E74_F*, and *E74_G* are active in salivary glands but inactive in wings during the larval-to-prepupal transition. Notably, *E74_D*, *E74_G*, and to a lesser extent, *E74_F* exhibit higher amplitude binding of EcR in salivary glands relative to wings, which correlates with greater activity of these enhancers in glands. This suggests that the relative levels of EcR occupancy between wings and salivary glands may predict differential enhancer activity; however, more examples of EcR-bound enhancers need to be tested, especially from targets besides primary response genes like *E74*. What is clear is that the absolute levels of EcR occupancy do not predict whether EcR is required for enhancer activity. We find that enhancers with high-amplitude EcR binding (e.g. *E74_A*) can be equivalently dependent on EcR for activation as enhancers with low-amplitude EcR binding (e.g. *E74_B*). Taken together, these findings indicate that ecdysone responses, even at primary response genes, are driven by a complex interplay between shared and tissue-specific enhancer activity.

One intriguing possibility is that regulation by multiple EcR-bound enhancers functions as a rheostat to fine-tune the duration or magnitude of ecdysone responses. It is also possible that additional layers of regulation exist downstream of EcR binding. Enhancers at primary response genes, including *E74*, have been shown to physically interact over long distances (86). This looping is dependent on nuclear pore complex members, and it promotes transcriptional memory to ecdysone exposure. Higher occupancy of EcR could facilitate looping between enhancers and promoters, leading to tissue-specific ecdysone responses for genes with differential EcR binding. Increased responsiveness of salivary glands to successive ecdysone pulses may also potentiate ecdysone responses in glands relative to wings. A role for NELF in re-induction of ecdysone-responsive genes has also recently been described, suggesting yet another level of regulation that may contribute tissue-specific ecdysone responses (87).

### Tissue-specific ecdysone responses are not driven by direct control of chromatin accessibility by EcR

The accessibility of cis-regulatory elements genome-wide plays an important role in establishing tissue-specific gene expression programs by influencing where transcription factors can bind (88-90). Transcription factors with the ability to control chromatin accessibility, collectively termed “pioneer factors”, have essential functions in both development and disease (91). Pioneer factors also perform key roles in shaping the transcriptional response to hormones (92), and nuclear receptors themselves have been found to possess pioneer-like activity (93). Given that EcR sits atop the genetic hierarchy that governs the ecdysone response, and due to the profound impact that ecdysone signaling has on gene expression (10, 28), we considered the possibility that EcR controls chromatin accessibility at its binding sites. In such a model, EcR binding at target enhancers would decrease nucleosome occupancy, rendering them competent to respond to tissue-specific transcription factors. EcR could perform this function either on its own or in collaboration with other transcription factors. EcR-dependent control of nucleosome occupancy could also involve nucleosome remodelers, which can increase transcription factor occupancy through sliding, remodeling, or eviction of nucleosomes to reveal DNA binding sites (94). Like other nuclear hormone receptors, ligand binding to EcR induces structural changes that result in differential association with co-repressor and co-activator complexes (95, 96), suggesting that control of chromatin accessibility could even be triggered by ecdysone pulses. Consistent with a potential role of EcR in controlling accessibility of target enhancers, examination of EcR DNA binding profiles revealed a strong association between tissue-specific EcR binding and tissue-specific open chromatin sites. However, EcR loss of function in the wing resulted in few changes in chromatin accessibility, and the changes that did occur are not associated with EcR binding. We conclude that EcR does not function as a binary switch to open chromatin at target enhancers.

The lack of large-scale changes in chromatin accessibility suggests that other transcription factors can still bind target enhancers in the absence of EcR. It is possible that EcR binding makes an additive contribution to chromatin accessibility at target enhancers because we detect a subtle decrease in chromatin accessibility at EcR binding sites in *EcR-RNAi* wings. It is also possible this decrease is due to loss of nucleosome remodelers recruited by EcR. Other assays besides FAIRE-seq that are better-suited for precise measurements of nucleosome positioning may provide further insight into chromatin changes surrounding EcR target enhancers. Regardless of the mechanism, loss of EcR function results in failure to activate target enhancers without producing dramatic effects on chromatin accessibility. Taken together, our findings indicate that tissue-specific responses to ecdysone are determined at least in part through binding of EcR to target enhancers made accessible by other transcription factors. Who might these transcription factors be? Remarkably, the ecdysone-induced transcription factor E93 is required for proper accessibility of temporally dynamic enhancers at later developmental stages (30). In the absence of E93, late-acting enhancers fail to open and activate, whereas early-acting enhancers fail to close and deactivate. This suggests a model wherein primary response factors specify temporal identity by determining target enhancer accessibility, and EcR contributes to this response by controlling the timing of enhancer activity as part of a feed forward loop. However, EcR does not determine which enhancers are available for use. In other words, despite its position at the top of the ecdysone-response cascade, EcR functions downstream of its own primary response genes in regulating target gene activity. Combined with the observation that primary responses to ecdysone vary considerably between tissues (**Figure 2**), we propose that a central step in regulating tissue-specific ecdysone responses lies in control over which primary response factors are induced, which in turn influences where EcR binds in the genome.

## Materials and Methods

### Immunofluorescence

Larvae were dissected in batches of 5-20 in PBS at room temperature and transferred to ice. Samples were then fixed in PBS plus 4% paraformaldehyde (Electron Microscopy Services) for 25-minutes at room temperature on an orbital shaker. Fixed samples were washed in PBS + 0.15% Triton X100 (PBT). Antibodies were incubated either for 2hrs at room temperature or overnight at 4°C. After secondary antibody incubation, samples were washed once with PBT, once with PBT + 0.2µg / ml DAPI, and once with PBT. The following antibody concentrations were used: 1:750 mouse anti-EcR (DSHB DDA2.7, concentrate), 1:4000 rabbit anti-GFP (Abcam ab290), 1:3500 mouse anti-Dl (DSHB C594.9b, concentrate), 1:200 mouse anti-FLAG M2 (Sigma F1804), 1:1000 anti-Br (DSHB 25E9.D7, concentrate). Secondary antibodies were: 1:1000 goat anti-rabbit, or goat anti-mouse, conjugated with either Alexa-488 or Alexa-594 (ThermoFisher A11037, A11034). Samples were imaged on a Leica Sp5 or Leica Sp8 confocal microscope.

### Sample preparation for RNAseq

A minimum of 60 wings or salivary glands were prepared as previously described (58) from either *Oregon R (WT)* or *yw; vg-GAL4, tub>CD2>GAL4, UAS-GFP, UAS-FLP / UAS-EcR-RNAi^104^ (EcR-RNAi)*. For library construction, 50-100ng RNA was used as input to the Tecan Genomics Universal RNA-Seq with NuQuant, Drosophila. Library preparation followed the manufacturer’s instructions with the following modifications: 1) after second-strand cDNA synthesis, samples were sonicated 5x20-seconds (30-seconds rest between cycles) on high power in a BioRupter bath sonicator; 2) qPCR was performed to determine the optimal cycle number using manufacturer’s recommendations; 3) after library amplification, an additional, 1.2:1 SPRI bead-cleanup was performed. Paired-end, 2x75 sequencing was performed on an Illumina HiSeq X using Novogene Co.

### Sample preparation for CUT&RUN

A minimum of 75 wings or 50 salivary glands from *w; EcR^GFSTF^/Df(2R)BSC313* were dissected in wash buffer (20mM HEPES-NaOH, 150mM NaCl, 2mM EDTA, 0.5mM Spermidine, 10mM PMSF). The rest of the protocol was performed as described in (97). Fragments that diffused out of the nucleus (“supernatant” sample), as well as size-selected DNA from the nuclei (“pellet” samples) were prepared and sequenced. For library preparation, the Takara ThruPLEX DNA-seq kit with unique dual-indexes was used following the manufacturer’s protocol until the amplification step. For amplification, after the addition of indexes, 16-21 cycles of 98C, 20s; 67C, 10s were run. A 1.2x SPRI bead cleanup was performed (Agencourt Ampure XP). Libraries were sequenced on an Illumina HiSeq 4000 with 2x75 reads. The following antibody concentrations were used: 1:300 mouse anti-FLAG M2; 1:200 rabbit anti-Mouse (Abcam ab46450); 1:400 Batch#6 protein A-MNase (gift of Steven Henikoff).

### Sample preparation for FAIREseq

Larvae from either *Oregon R (WT)* or *yw; vg-GAL4, tub>CD2>GAL4, UAS-GFP, UAS-FLP / UAS-EcR-RNAi^104^ (EcR-RNAi)* were dissected in 1xPBS in batches of 5-10 then fixed at RT for 10-minutes in 4% paraformaldehyde, 50mM HEPES (pH 8.0), 100mM NaCl, 1mM EDTA (pH 8.0), 0.5mM EGTA (pH 8.0). Fixation was quenched by incubation for 5m in 1xPBS, 125mM Glycine, 0.01% Triton X-100 and then transferred to 10mM HEPES (pH 8.0), 10mM EDTA (pH 8.0), 0.5mM EGTA (pH 8.0), 0.25% Triton X-100, 1mM PMSF. Wings or salivary glands were dissected off cuticles and snap frozen in liquid nitrogen. Samples were lysed in 2% Triton X-100, 1% SDS, 100mM NaCl, 10mM Tris (pH 8.0), 1mM EDTA. Following lysis, a minimum of 40 wings or salivary glands were pooled together and homogenized using 2.38mm tungsten beads with 6 cycles of 1min on and 2min off and then sonicated using a Branson Sonifier with 5 cycles of 30-seconds (1-second on, 0.5-second off) while letting the samples rest for at least 2-minutes on ice between cycles. An aliquot was removed as an input fraction. The remaining samples were subjected to phenol-chloroform and chloroform extractions and then precipitated with ethanol. Input and experimental samples were heated overnight at 65C to reverse cross links and then treated with RNase A for 1-hour at 37°C. DNA was purified with a Qiagen QIAquick PCR Purification Kit eluting in nuclease free water. Samples were used as input into the Takara ThruPLEX DNA-seq kit following manufacturer’s instructions.

### RNA Sequencing Analysis

Reads were trimmed using bbmap (v38.75) with parameters ktrim=r ref=adapters rcomp=t tpe=t tbo=t hdist=1 mink=11. Reads were aligned with STAR (2.7.3a) (98). Indexes for STAR were generated with parameter --sjdbOverhang 74 using genome files for the dm6 reference genome. The STAR aligner was run with parameters --alignIntronMax 50000 --alignMatesGapMax 50000. Samtools (v1.9) was used to filter reads to those with a q-score greater than 2. RSubread (v2.0.1) was used to count reads mapping to genes using a gtf file from flybase.org (r6.32) using parameters: annot.ext = gtfPath, isGTFAnnotationFile = T, isPairedEnd = T, strandSpecific = 1, nthreads = 4, GTF.featureType = ’exon’, allowMultiOverlap = F (99). DESeq2 (v1.26.0) was used to identify differentially expressed genes using the lfcShrink function to shrink log-fold changes and with each genotype and time-point as a separate contrast (100). Differentially expressed genes were defined as genes with an adjusted p-value less than 0.05 and an absolute log_2_ fold change greater than 1. Normalized counts were generated using the counts function in DESeq2. For c-means clustering, normalized counts were first converted into the fraction of maximum normalized counts across all tissues and conditions and c-means clustering was performed using the ppclust package (v1.1.0) (101). MA Plots were made with ggplot2 and points were shaded using kernel density estimates calculated using the MASS (v7.3-51.4) package (102). Heatmaps were generated using ggplot2 (v3.3.2) and patchwork (v1.1.0) in R (103-105). Gene Ontology (GO) analysis was performed using Bioconductor packages TopGO (v2.38.1) and GenomicFeatures (v1.38.2) using expressed genes as a background set with parameters: algorithm = ‘elim’ and statistic = ‘fisher’ (106). Similar GO terms were collapsed based on semantic similarity using the rrvgo package in R and only the parent term was used (v1.1.1) (107). Expressed genes were defined as genes with a normalized count value >= 10.

### CUT&RUN Sequencing Analysis

Technical replicates were merged by concatenating fastq files. Reads were trimmed using bbmap (v38.75) with parameters ktrim=r ref=adapters rcomp=t tpe=t tbo=t hdist=1 mink=11. Trimmed reads were aligned to the dm6 reference genome using Bowtie2 (v2.2.8) with parameters --local --very-sensitive-local --no-unal --no-mixed --no-discordant --phred33 -I 10 -X 700 (108). Reads with a quality score less than 5 were removed with samtools (v1.9) (109). PCR duplicates were marked with Picard (v2.21) and then removed with samtools. Fragments between 20 and 120bp were isolated using a custom awk script and used for downstream analyses as recommended in (110). Bam files were converted to bed files with bedtools (v2.29) with parameter -bedpe (111). Bedgraphs were generated with bedtools and then converted into bigwigs with ucsctools (v320) (112). Data was z-normalized using a custom R script. MACS (v2.1.2) was used to call peaks on individual replicates and merged files using parameters -g 137547960--nomodel --seed 123 (113). As a control for peak calling, wing IgG supernatant and pellet samples were used. Wing IgG controls were *yw* CUT&RUN samples in which the primary antibody was omitted and only the mouse anti-Rabbit IgG secondary was used. To identify differentially bound regions, a union peak set was generated and RSubread (v2.0.1) was used to assign to features using parameters strandSpecific = 0, allowMultiOverlap = T and then used as input for DESeq2 (v1.26.0) (114, 115). To identify sites that were differentially bound in each tissue irrespective of whether they were found in the supernatant or pellet samples (see Sample Preparation for CUT&RUN), we entered the DESeq2 design formula as, “∼tissue + supPel”. For pairwise comparisons, union peaks were subsequently filtered to contain peaks that overlapped a peak found in either sample by at least one base pair. MA plots were made as described for RNAseq. Heatmaps and average signal plots were generated from z-normalized data using the Bioconductor package Seqplots (v1.24.0) and plotted using ggplot2 (116). ChIPpeakAnno (v3.20.0) was used to calculate distance of peaks to their nearest gene (117). To identify clusters of EcR binding sites, the EcR peaks were resized to 5000bp, assigned to clusters, and the furthest start and end coordinate of the original peaks were used.

### FAIRE sequencing analysis

Technical replicates were merged by concatenating fastq files. Reads were trimmed using bbmap (v38.75) with parameters ktrim=r ref=adapters rcomp=t tpe=t tbo=t hdist=1 mink=11. Trimmed reads were aligned to the dm6 reference genome using Bowtie2 (v2.2.8) with parameters --phred33 --seed 123-x (118). Reads with a quality score less than 5 were removed with samtools (v1.9) (119). PCR duplicates were marked with Picard (v2.21) and then removed with samtools. The remaining processing and analysis steps were performed as described for CUT&RUN.

### Motif Analysis

To identify occurrences of the EcR motif in the genome, PWMs for the EcR and Usp motifs identified by a bacterial 1-hybrid were obtained from Fly Factor Survey (120). For the palindromic, Usp/EcR motif, the PWMs for EcR and Usp were concatenated together and the probabilities for the central, overlapping base were averaged. FIMO (v4.12.0) was run on the dm6 reference genome using parameters –max-stored-scores 10000000 --max-strand --no-qvalue --parse-genomic-coord --verbosity 4 --thresh 0.01 (121). Motif density plots were generated by counting the number of motifs from peak summits (10bp bins) and normalizing by the number of input peaks.

### EcR knockdown in the wing and salivary gland

To knockdown EcR in the wing and salivary gland in parallel, we made use of the previously published line: *yw; vg-GAL4, UAS-FLP, Tub>>STOP>>GAL4, UAS-GFP / CyO* (122). Early activation of *vg-GAL4* throughout the wing primordia results in flip-out of the stop-cassette and persistent expression of *Tub-GAL4* throughout wing development. This construct is also active in the salivary gland, which may be a consequence of the *vg-GAL4* p-element vector which has been previously reported to have a minimal promoter active in the salivary gland (123-126).

### Drosophila culture and genetics

Flies were grown at 25C under standard culture conditions. Late wandering larvae were used as the – 6hAPF timepoint. White prepupae were used as the 0h time point for staging +6hAPF animals. For staging –30hAPF, apple juice plates with embryos were first cleared of any larvae. Four hours later, any animals that had hatched were transferred to vials. 72 hours later, tissues were harvested. The following genotypes were used: *yw; vg-GAL4, UAS-FLP, UAS-GFP, Tub>CD2>GAL4 / CyO* (122)*. W1118; P{UAS-EcR-RNAi}104 (BDSC#9327) yw; EcR^GFSTF^ (BDSC#59823) w1118; Df(2R)BSC313 /CyO (BDSC#32253) yw; + / +; br^disc^::tdTomato / TM6B yw; +/+; E74_A-I::tdTomato/TM6B*

---

**Figure S1.**
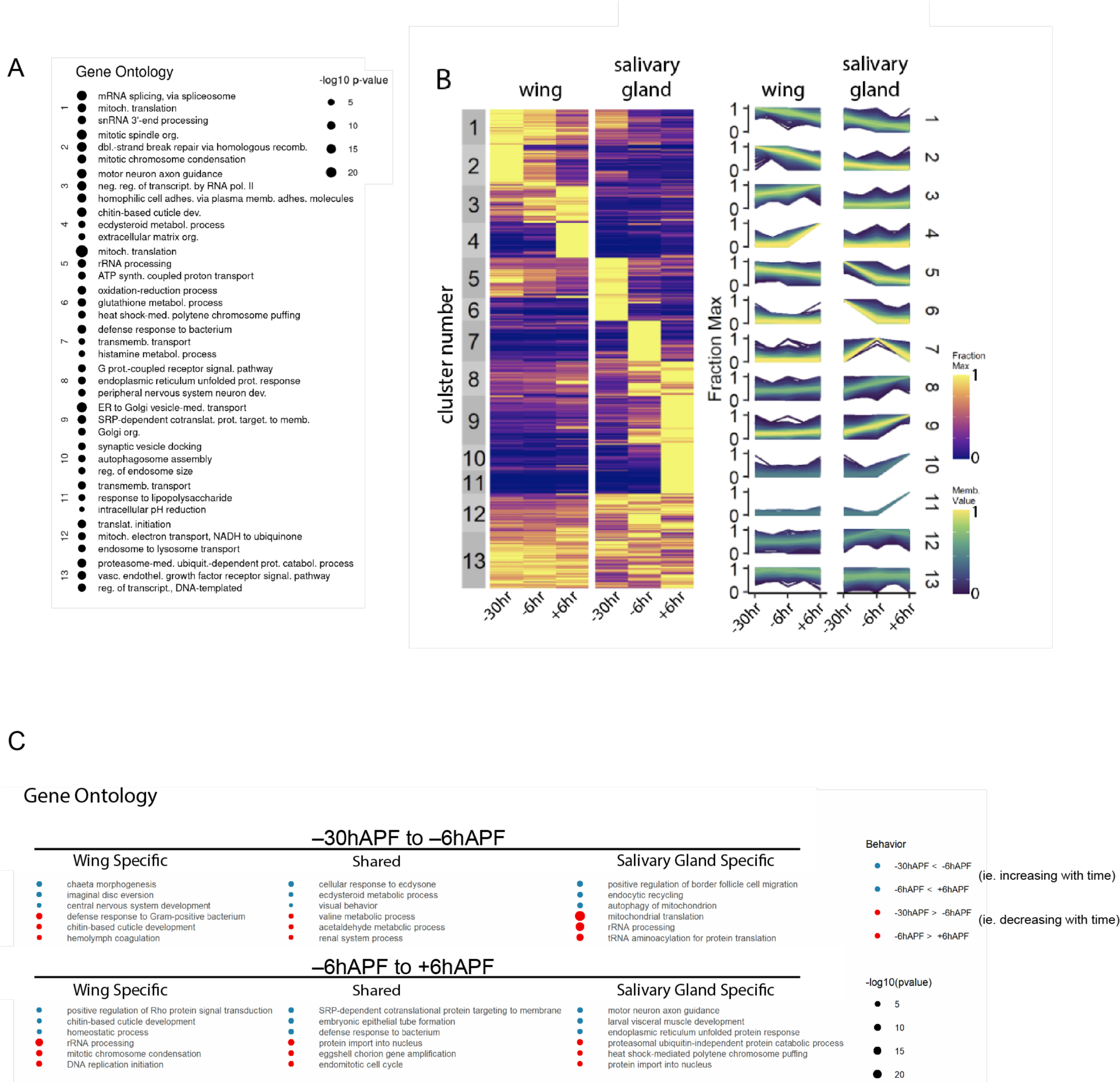
Gene ontology of temporally dynamic genes in wings and salivary glands. (A) Gene ontology terms for each cluster of genes depicted in the heatmap shown in Figure 1B. (B) Copy of the heatmap RNA-seq clusters shown in Figure 1B. (C) Gene ontology terms for temporally dynamic genes between adjacent time points in wild-type wings and salivary glands; these are the same genes depicted in the Venn diagram shown in Figure 1C.

**Figure S2.**
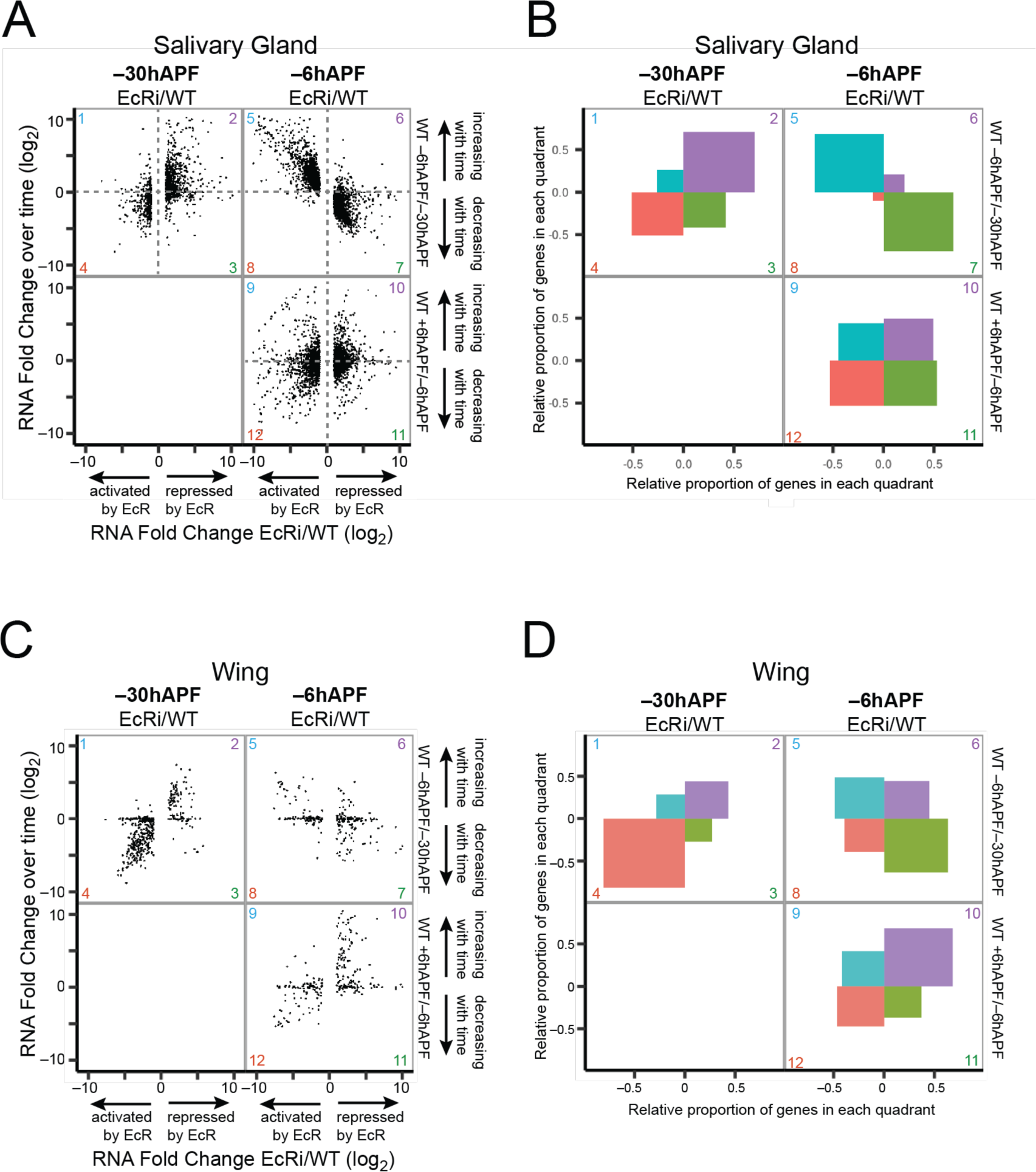
EcR is required for both gene activation and gene repression at –30hAPF and at –6hAPF in wings and salivary glands. (A) and (B) Copies of the scatterplots and gene proportion plots from Figure 2C, D. (C) Scatterplots of RNA-seq values for differentially expressed genes in *EcR-RNAi* wings. The ratio between *EcR-RNAi* and wild-type is shown on the x-axis for –30hAPF and –6hAPF. The ratio between adjacent wild-type stages is shown on the y-axis. (D) Plots indicating the proportion of genes located in each quadrant for the three scatterplots shown in Panel C.

**Figure S3.**
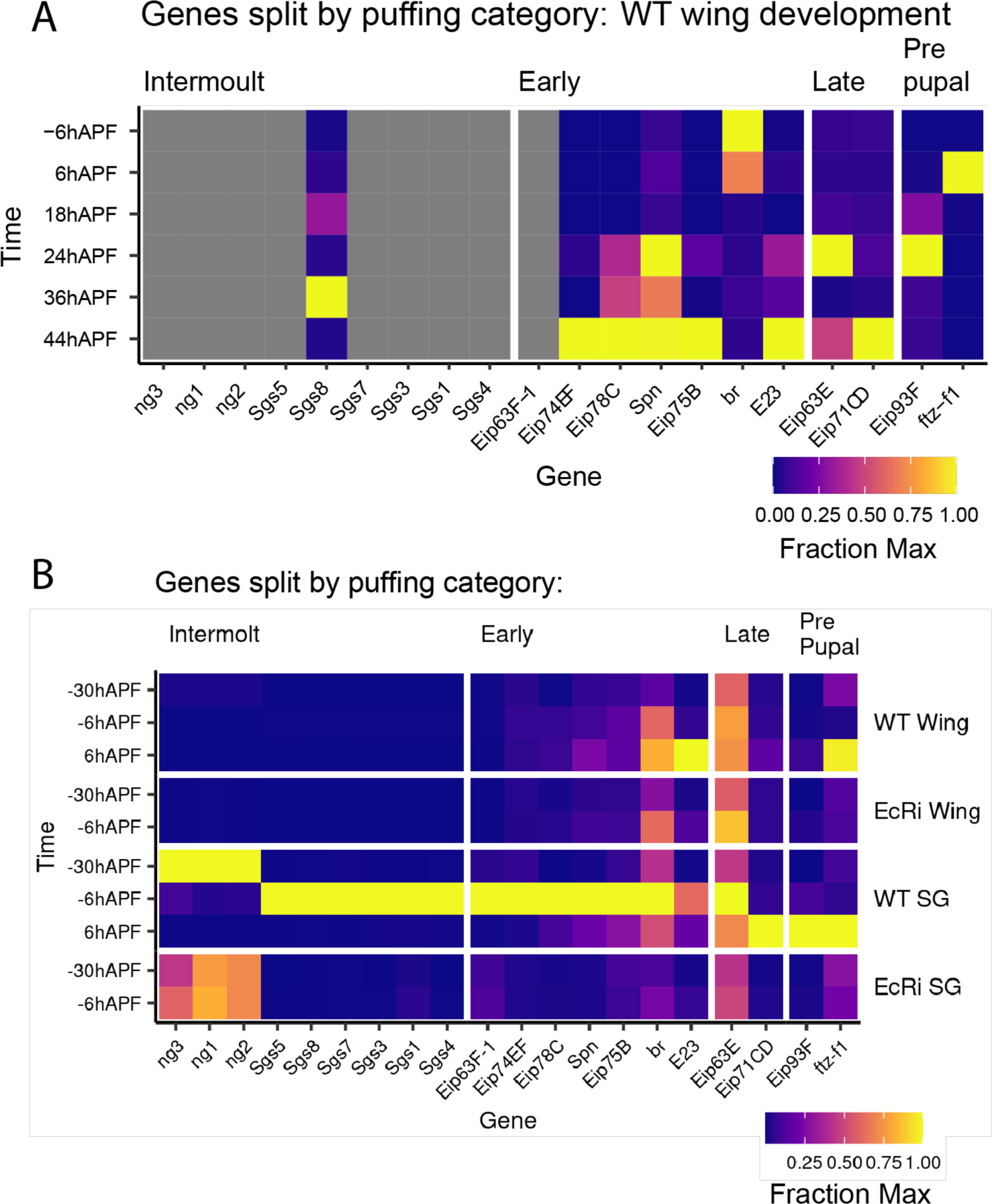
Expression of genes residing in puffing loci from developing wings. (A) Heatmap of RNA-seq values (fraction of max) from a wild-type wing developmental time course for select genes that exhibit ecdysone-dependent puffs in the salivary gland. (B) Heatmap of RNA-seq values (fraction of max) from wild-type and *EcR-RNAi* tissues for select genes that exhibit ecdysone-dependent puffs in salivary glands.

**Figure S4.**
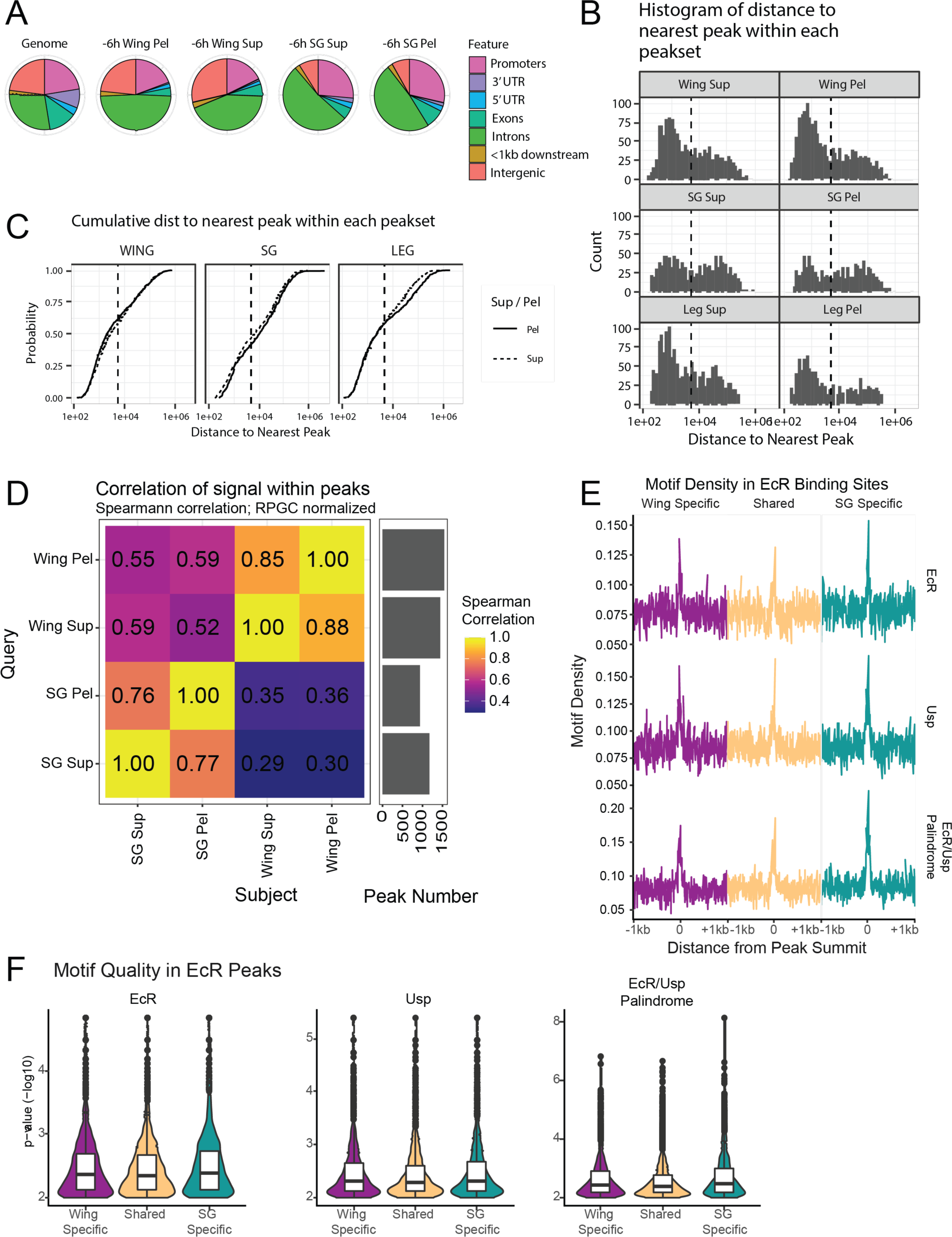
Additional properties of EcR binding sites. (A) Pie charts of the genome-wide distribution of EcR binding sites. (B) Histograms and (C) cumulative distribution plots of the distance of each EcR peak to its nearest neighbor. Peaks falling within 5kb (dotted lines) of each other define an EcR cluster. (D) Heatmap of the Spearman correlation coefficient for EcR CUT&RUN signal within EcR peaks between tissues and separated by whether the supernatant (sup) or pellet (pel) DNA was sequenced following the CUT&RUN protocol (see Methods). (E) Line plots of motif density surrounding the summit (+/-1kb) of EcR CUT&RUN peaks. Individual EcR and Usp motifs from Fly Factor Survey were used, as well as an EcR/Usp palindrome that was generated by combining EcR and Usp motifs. (F) Violin plots of p values for sequences matching EcR, Usp individual motifs, or the EcR/Usp palindrome.

**Figure S5.**
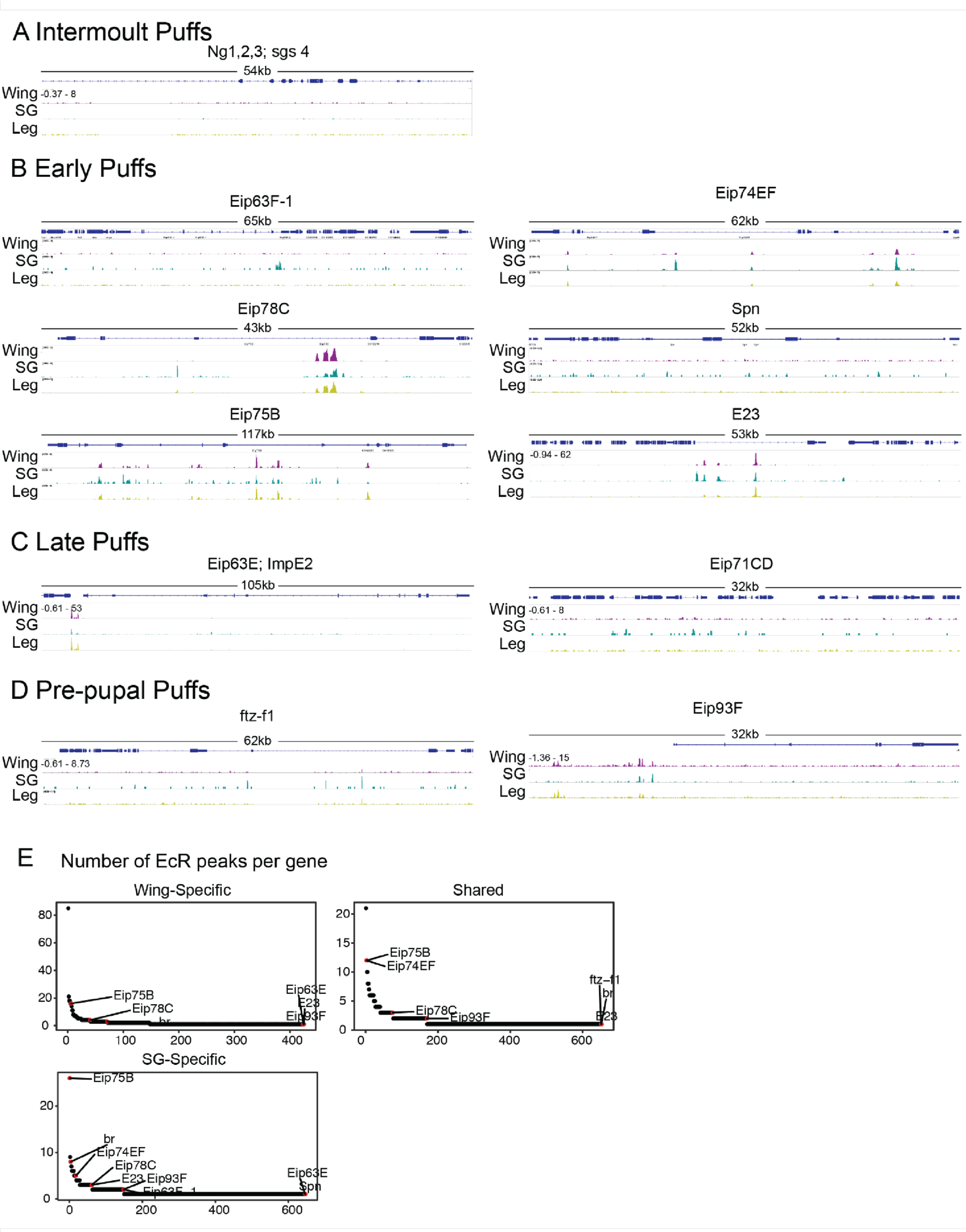
EcR binds extensively near ecdysone primary response genes using a mixture of tissue-specific and shared binding sites. (A-D) Browser shots of EcR CUT&RUN signal in wings and salivary glands at puffing loci. (E) Ranked plots of EcR peak number at wing-specific, shared, and salivary gland-specific EcR CUT&RUN peaks. Puffing genes are indicated and highlighted in red.

**Figure S6.**
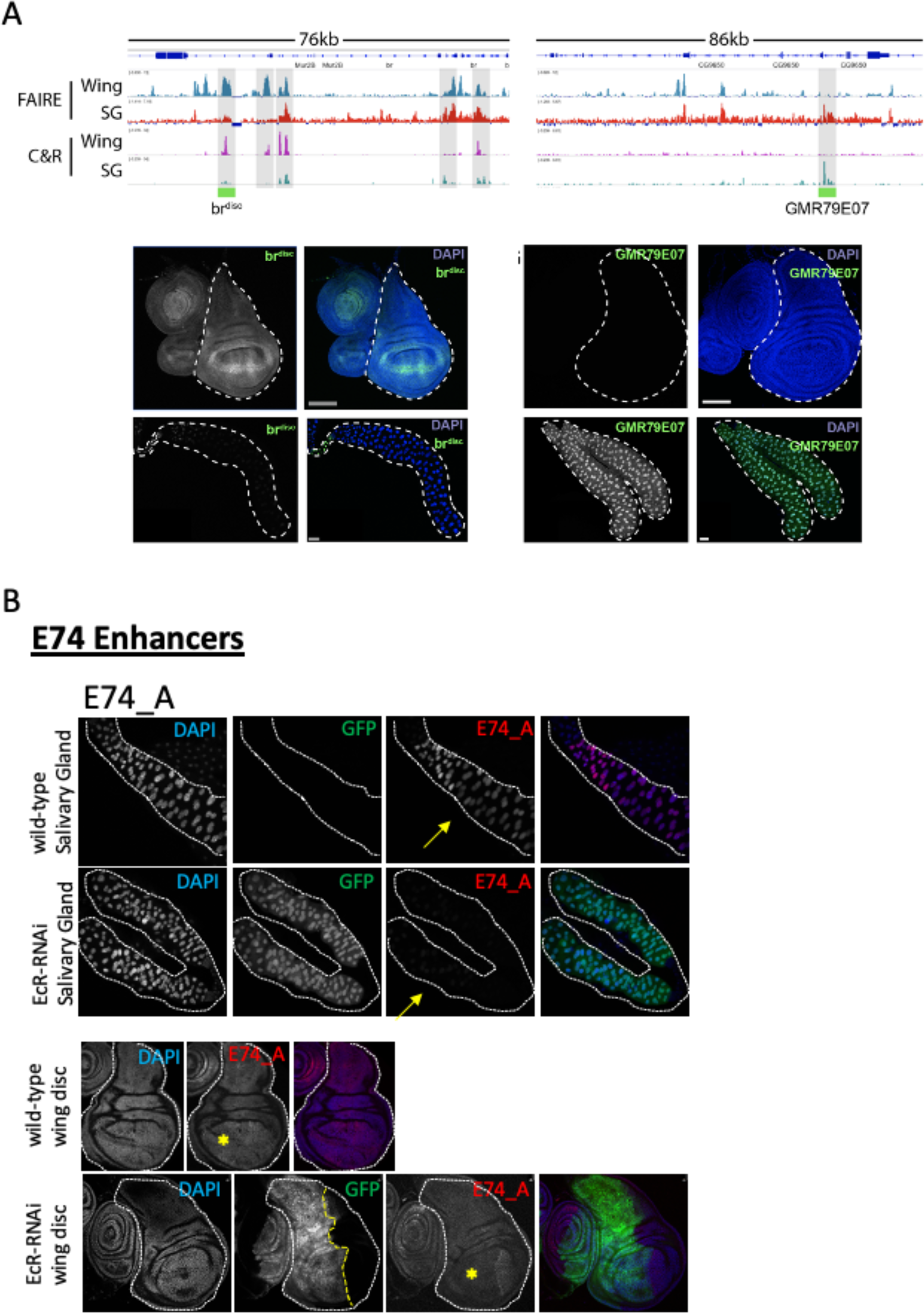
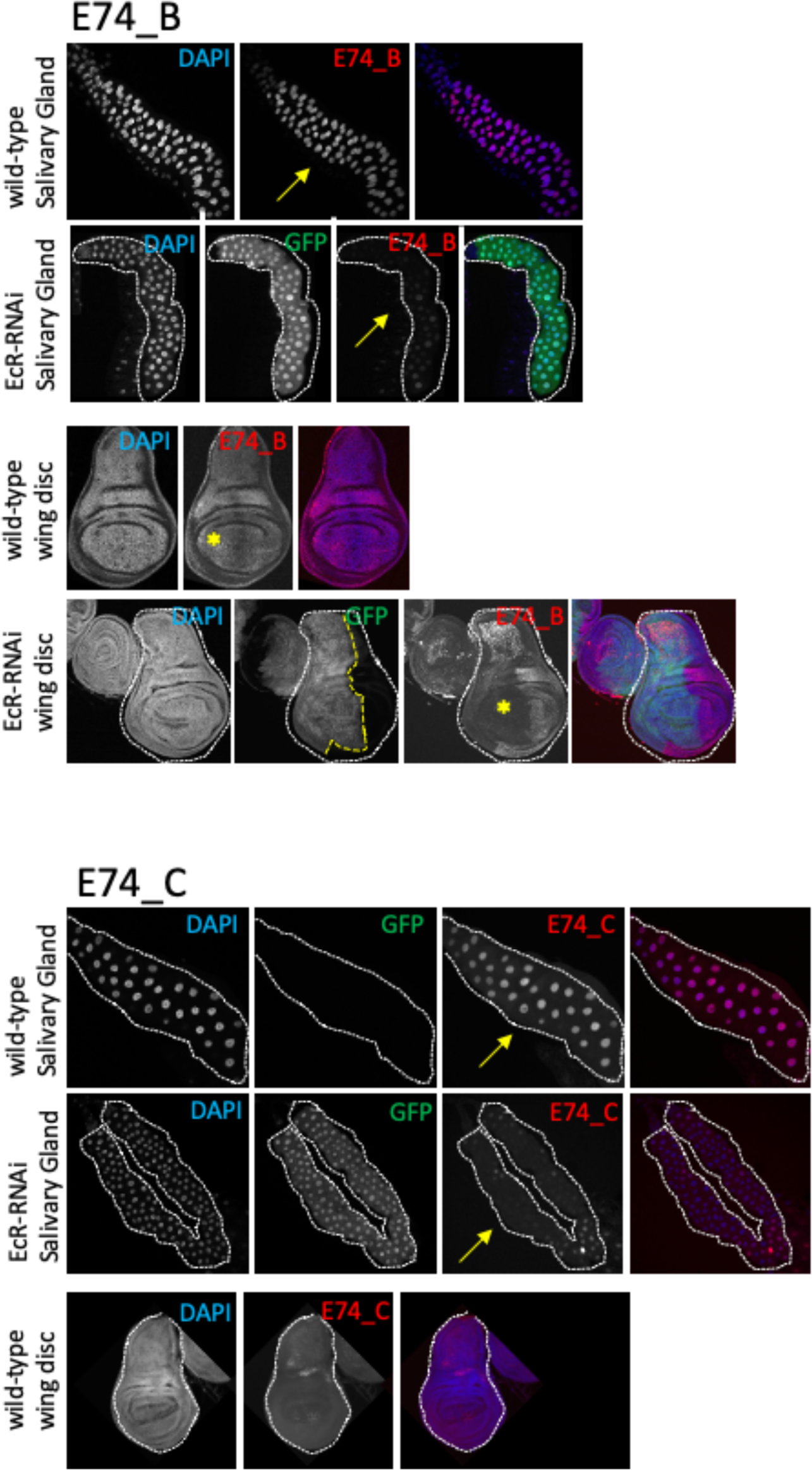
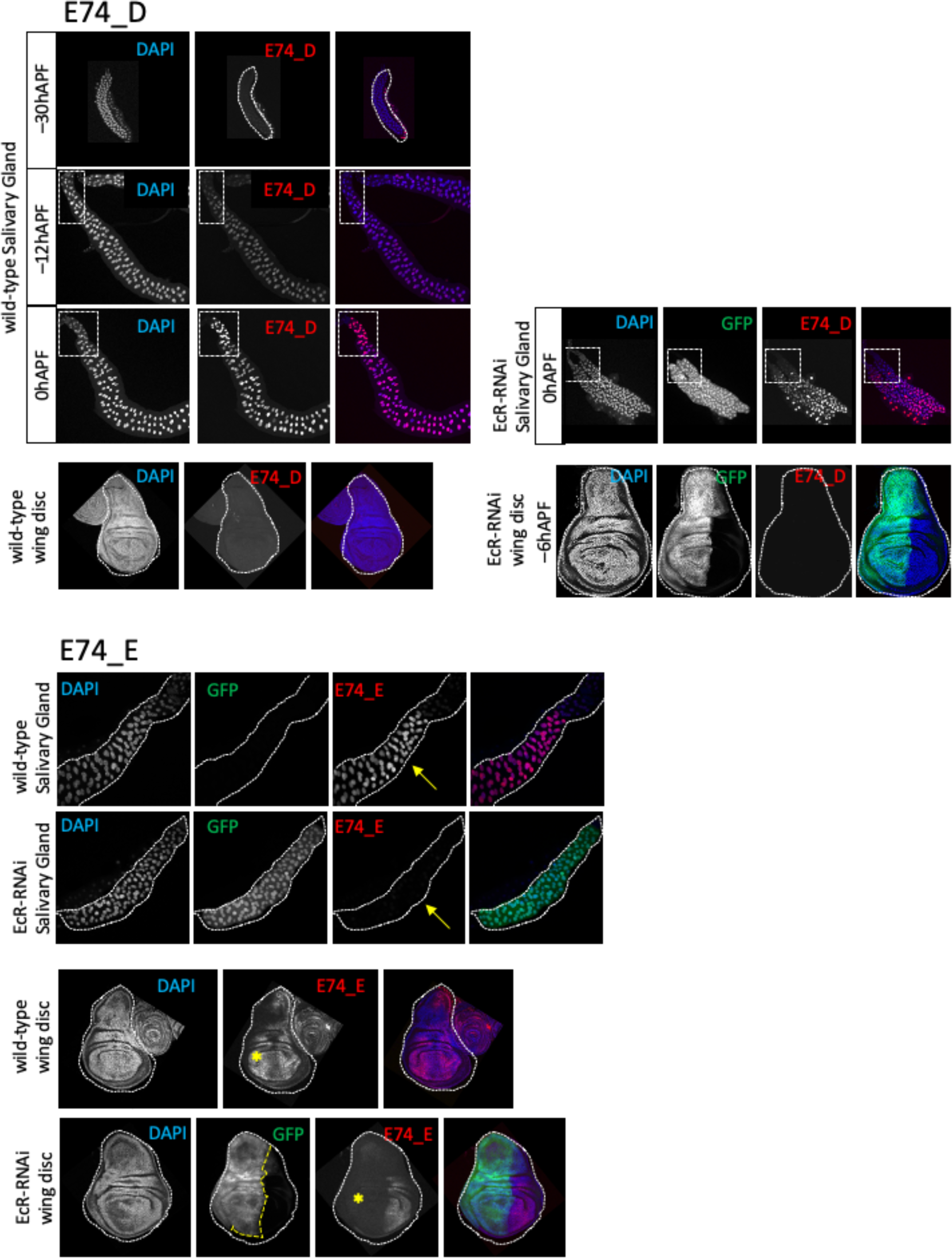
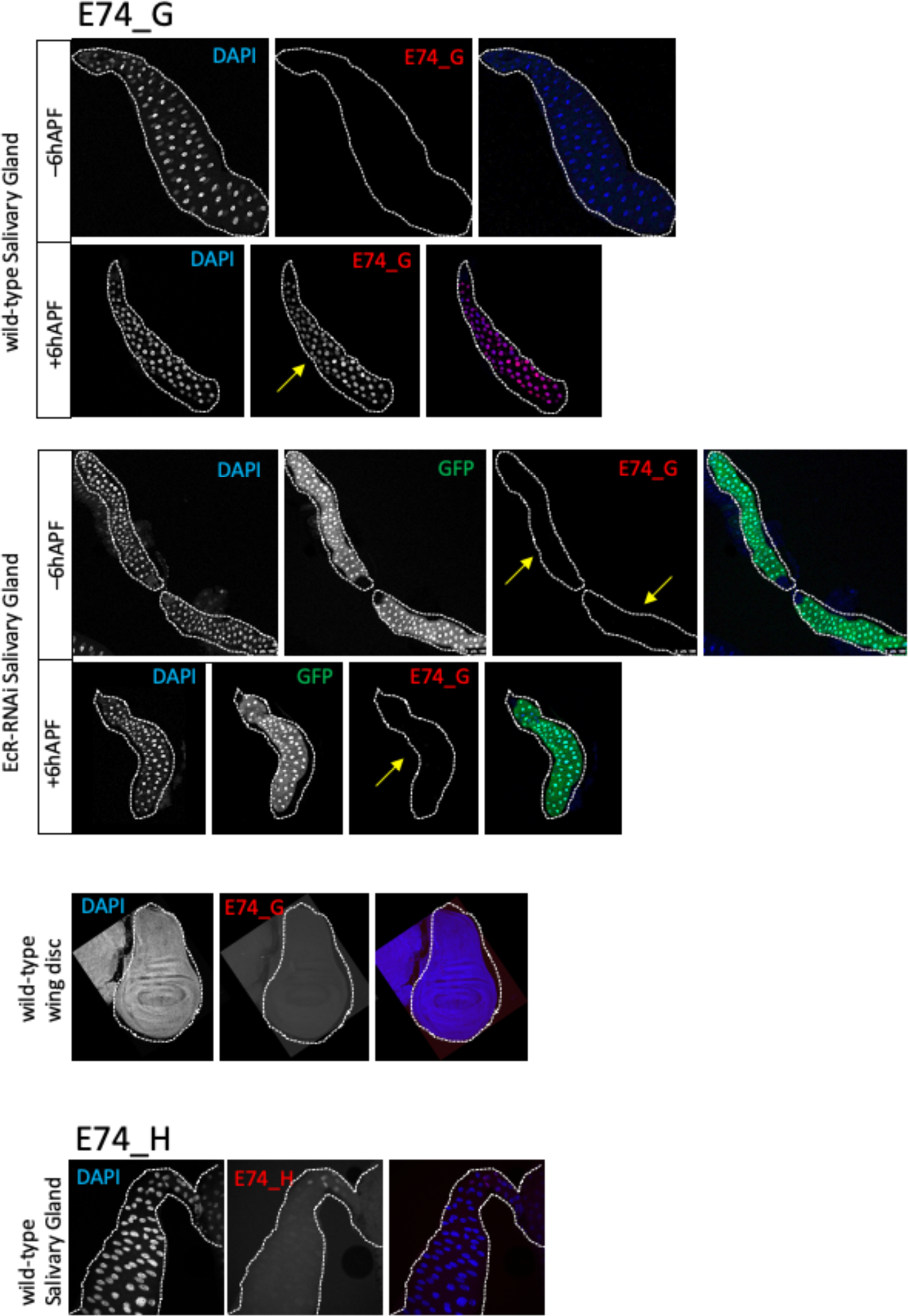
EcR binding sites correspond to active enhancers. (A) Browser shots of FAIRE and EcR CUT&RUN signal from –6hAPF wings and salivary glands for genomic loci surrounding the *br^disc^* and *GMR79E07* enhancers. Confocal images depict enhancer activity (green) and DAPI (blue) in wild-type wings and salivary glands from the same developmental stage. (B) Confocal images of *E74* enhancer activity in wild-type wings and salivary glands. Enhancer activity is visualized by tdTomato expression (red), DNA by DAPI stain (blue). For *EcR-RNAi* tissues, the expression of the GAL4 driving *UAS-EcR-RNAi* is captured by *UAS-GFP* (green). Yellow lines in wing discs indicate the boundary between *Ci-GAL4* expressing and wild-type cells. Developmental stage is –6hAPF for all images, except when indicated on the left. Yellow asterisks indicate loss of enhancer activity relative to wild-type wings. Yellow arrows indicate loss of enhancer activity relative to wild-type salivary glands. For *E74_D*, the white dashed box highlights the transition cells, which exhibit the strongest dependence on EcR for enhancer activity. In wild-type glands, *E74_D* enhancer activity increases in transition cells during the larval-to-prepupal transition; however, in *EcR-RNAi* cells, the enhancer fails to increase in activity in these cells.

**Figure S7.**
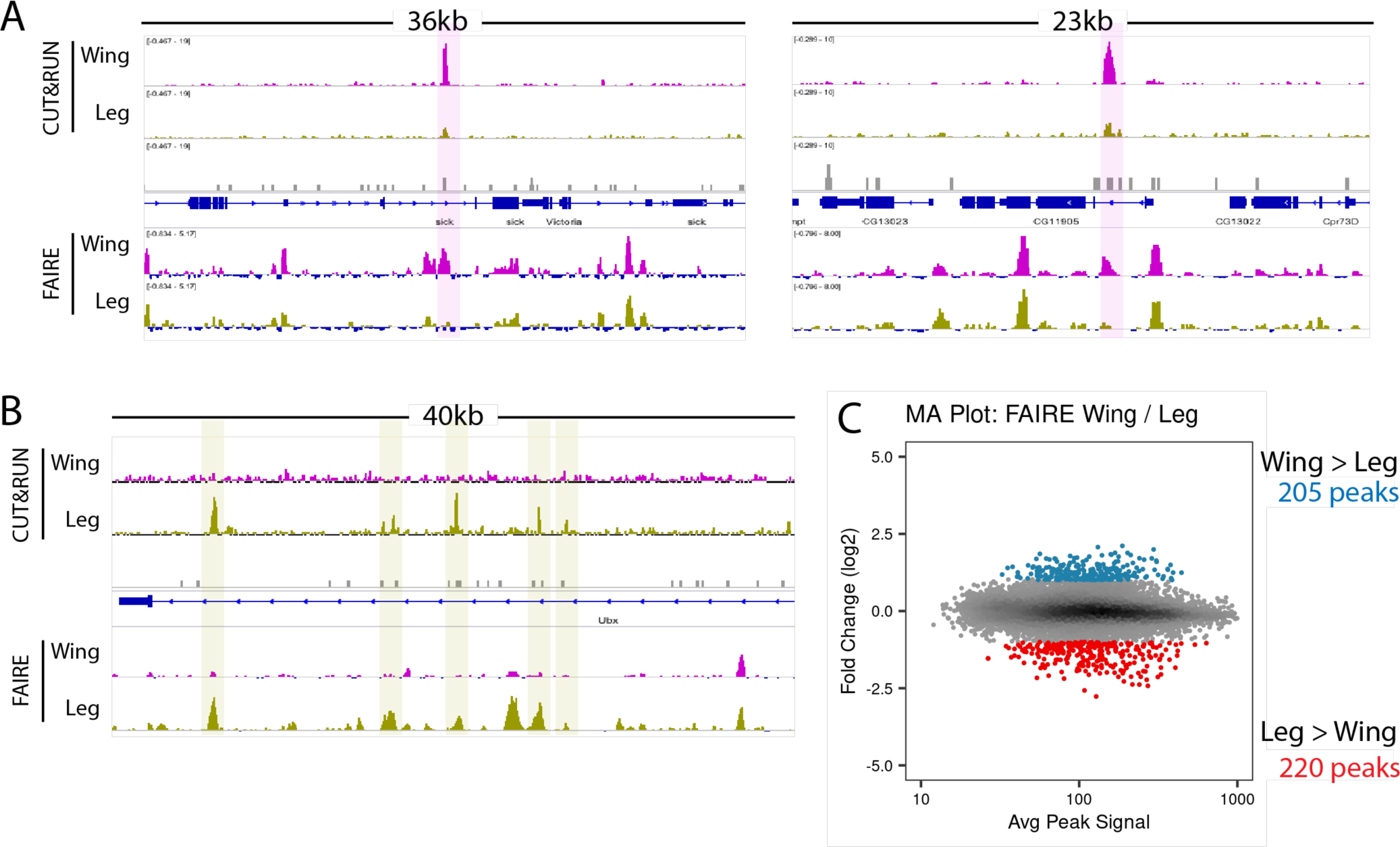
Differential EcR binding between wings and legs overlaps sites of differential accessibility. (A) Browser shots of EcR CUT&RUN and FAIRE signal from –6hAPF wings and legs at loci that exhibit wing-specific EcR peaks. (B) Browser shots of EcR CUT&RUN and FAIRE signal from wings and legs at loci that exhibit leg-specific EcR peaks. (C) MA plot of FAIRE signal in –6hAPF wings and legs. Differential peaks (absolute log_2_ fold change > 1, adj p value < 0.05) are colored in blue and red.

## Notes

### Competing Interest Statement

The authors have declared no competing interest.

